# Protein kinase CK2α’ as a dual modulator of immune signaling and synaptic dysfunction in Tauopathy

**DOI:** 10.1101/2025.06.24.661391

**Authors:** Angel White, Peter Gavrilyuk, Persephone Gu, Rafael Falcon Moya, Reid Thurston, Amal Fickak, Nicholas B Rozema, Prarthana Keshavaram, Scott Vermilyea, Riley Schlichte, Joyce Meints, Ying Zhang, Alfonso Araque, Michael Lee, Rocio Gomez-Pastor

**Affiliations:** University of Minnesota, School of Medicine, Department of Neuroscience, Minneapolis, MN, USA.; Minnesota Supercomputing Institute, University of Minnesota, Minneapolis, MN, USA

## Abstract

Tauopathies are a group of neurodegenerative diseases characterized by tau accumulation, neuroinflammation, and synaptic dysfunction, yet effective treatments remain elusive. Protein Kinase CK2 has been previously associated with different aspects of tau pathology but genetic evidence for the contribution of CK2 to tauopathy remained lacking. Here, we show CK2α’, one of the two catalytic subunits of CK2, as a novel regulator of tau-mediated neurodegeneration. We found that CK2α’ expression is elevated in postmortem brains of dementia patients and in the hippocampus of PS19 tauopathy mice, especially in neurons and microglia. Using genetic haploinsufficiency in PS19 mice, we demonstrated that reduced CK2α’ levels significantly decrease phosphorylated tau and total tau burden in the hippocampus and cortex. CK2α’ depletion also attenuated microglial activation, pro-inflammatory cytokine production, and microglia synaptic engulfment, enhanced synaptic gene expression, synaptic density, and LTP. Importantly, CK2α’ depletion rescued cognitive deficits assessed in the Barnes maze. These effects appear to be mediated through both neuronal and glial functions and may involve CK2α’-dependent modulation of tau-associated phosphorylation and neuroinflammatory and immune signaling pathways.

**One Sentence summary:** This study highlights CK2α’ as a key node at the intersection of tau pathology, synaptic dysfunction, and neuroimmune signaling.

## Background

Tauopathies encompass a broad range of neurodegenerative conditions, most notably Alzheimer’s disease (AD) and related dementias (ADRD), for which no cure currently exists (*1–3*). Tauopathies are typically marked by progressive neuronal loss, synaptic and cognitive dysfunction, and widespread inflammation in both the central nervous system and peripheral tissues(*4–8*). A deeper understanding of the molecular mechanisms underlying tauopathies and their downstream consequences is essential for developing effective disease-modifying therapies. One key area of research is the role neuroinflammation plays in the progression of tauopathies. These disorders are characterized by chronic, excessive inflammatory responses to abnormal tau proteins (*5, 6, 9*). This persistent inflammation can exacerbate tau pathology by promoting sustained neuronal damage and dysfunction (*9–12*). However, despite ongoing research, the mechanisms that connect neuroinflammation and tau pathology are still poorly understood.

Protein kinase CK2 (formerly known as Casein Kinase II), is a serine/threonine kinase that has emerged as an important regulator of neuroinflammation and protein homeostasis in several neurodegenerative diseases (*13–15*), and it has previously been connected to tau pathology in AD (*16–19*). However, its specific role in the inflammatory pathways associated with tauopathies remains poorly defined. Interestingly, CK2 has demonstrated potential as a therapeutic target in various neurodegenerative contexts (*14, 15*). In AD, CK2 has been implicated in processes such as synaptic plasticity (*20, 21*), amyloid precursor protein (APP) processing (*22–24*), and tau accumulation (*18, 19*), highlighting its relevance for further investigation.

CK2 is a tetrameric enzyme composed of 2 regulatory β subunits (CK2β) and 2 catalytic subunits α (CKα) or α’ (CK2α’) (*13*). CK2α and CK2α’ are highly similar in structure but display differential expression and substrate specificity (*25–31*). CK2α is expressed throughout the body at relatively equal levels and has hundreds of substrates, while CK2α’ expression is more restricted to testes and brain and has very few validated substrates (*25–31*). Another important difference between these two catalytic subunits is the embryonic lethality of CK2α knock-out mice, while complete deletion of CK2α’ does not have any major defects other than male sterility (*30, 31*). Recent studies show that CK2α’ is selectively upregulated in HD, and partial genetic reduction of CK2α’ in HD mouse models improves synaptic function, reduces protein aggregation, and alleviates behavioral deficits, indicating its key role in disease pathology (*14, 15*).

CK2 was one of the first protein kinases identified to be altered in an AD brain (*32, 33*), but its role in AD has been somewhat controversial due to discrepancies in the ability to detect consistent alterations in different affected tissues. Early studies using pan-CK2 antibodies (non-selective for CK2α’/CK2α subunits) showed increased levels of total CK2 in the hippocampus and frontal cortex of AD mouse models and patients with AD, especially in glial cells (*34*) but others have reported opposite results (*32, 35, 36*). In those studies where levels of CK2 were elevated, the authors associated the increase in CK2 immunoreactivity with the activation of neuroinflammatory processes and AD pathology (*34*).

Although genetic studies have not yet clarified whether CK2α and CK2α’ differentially contribute to AD pathology, multiple lines of evidence implicate CK2 in Tau phosphorylation and tau-mediated dysfunction. Kinase inhibitor screens identified CK2 as a major driver of okadaic-acid–induced pTau in N2a cells (*17*), and CK2-mediated phosphorylation of the PP2A inhibitor SET increases pTau levels in neurons exposed to Aβ or overexpressing hTau (*16*). Moreover, mice overexpressing CK2 exhibit elevated pTau, impaired synaptic plasticity, and cognitive deficits reminiscent of tauopathies (*16*). Despite this, most studies have treated CK2 as a single holoenzyme, overlooking known differences in substrate specificity between CK2α and CK2α’. Determining whether these catalytic subunits play distinct roles in tauopathy is essential for developing targeted therapies, especially given the current lack of subunit-selective CK2 inhibitors and the opportunity this presents for designing more precise and effective treatments.

Due to previous findings specifically connecting the dysregulation of CK2α’ with various neurodegenerative diseases and reports linking this subunit to the phosphorylation of Tau in different contexts, we explored the role of CK2α’ in various tauopathy models. We confirmed a specific upregulation at the RNA and protein levels of CK2α’ in affected tissues of AD and FTD patients as well as a mouse model of tauopathy (PS19). CK2α’ upregulation was specifically observed in both neurons and microglia. Importantly, genetic depletion of CK2α’ in both cell and mouse models of tauopathy (expressing Tau-P301L or Tau-P301S mutations) resulted in decreased pTau. Furthermore, PS19 mice haploinsufficient for CK2α’ showed attenuated microglial activation, pro-inflammatory cytokine production, and microglia synaptic engulfment, enhanced synaptic gene expression, synaptic density, and LTP. Importantly, CK2α’ depletion rescued cognitive deficits assessed in the Barnes maze. Overall, our data showed a large role of the CK2α’ catalytic subunit specifically in tau pathology, largely connected to the modulation of microglia and synaptic functions. These studies highlight CK2α’ as an attractive therapeutic target for the treatment of AD/ADRD.

## Results

### Levels of Protein kinase CK2α’ are increased in the brains of dementia patients and mouse models of tauopathy

There are two different genes (*Csnk2a1* and *Csnk2a2*) that code for two different CK2 catalytic subunits CK2α and CK2α’, respectively. These two catalytic subunits share a high percent of homology and they can arrange in various combinations (α-α, α-α’ or α’-α’) depending on their expression and tissue availability although the canonical arrangement in the brain is expected to be α-α’ (**Fig. 1A, B**) (*13, 25–31*). Analyzing existing RNA-seq data from the Allen Brain Institute: Aging, Dementia, and Traumatic Brain Injury (TBI) Study (*37*) we found a specific increase in the expression of *CSNK2a2* (CK2α’), but not in *CSNK2a1* (CK2α), in the parietal cortex (PCx) of patients with dementia compared to non-dementia controls (**Fig. 1C-E**). We also found a trend towards increased CK2α’ in the temporal cortex (TCx) and hippocampus (HIP) although data did not reach statistical significance (**Fig. 1C**) (*37*). Importantly, we also observed a significant increase in CK2α’ protein levels, but not CK2α, in the frontal cortex of FTD patients compared to age and sex-match controls (**Fig.1F left panel, Fig. 1 G, H, File S1**). Immunoblotting for CK2α’ in CK2α’ knock-out mice confirmed antibody specificity (**Fig. 1F, right panel**). Enhanced levels of CK2α’ was also associated with the presence of tau pathology, as previously assessed by AT8 immunostaining and reported in the sample metadata provided by the NeuroBioBank (**Fig. 1I, File S1**). The majority of FTD samples grouped in the top left quadrant representing high levels of CK2α’ and the presence of tau pathology, while most control samples grouped in the bottom right quadrant representing low levels of CK2α’ and absence of tau pathology. For those two control samples (#1 and #6) in which tau pathology was observed, the AT8 phospho-tau (pTau) signal and distribution differed from the canonical tau pathology associated with dementia and individuals lacked classical cognitive decline (**File S1**).

**Figure 1.**
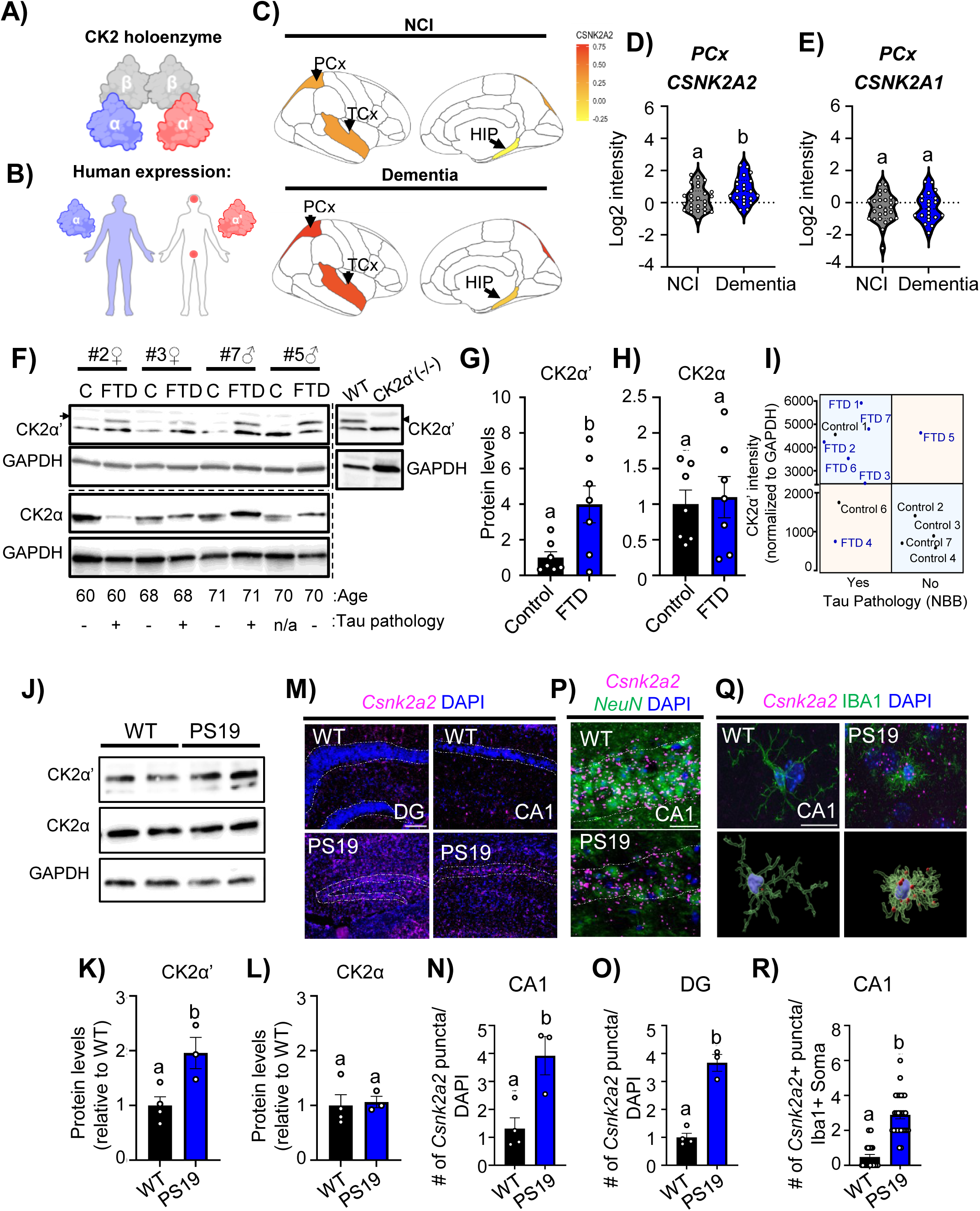
CK2α’ is increased in tissues of patients with AD/ADRD and in PS19 mice. **(A, B)** Representative diagram of CK2 holoenzyme, including regulatory and catalytic subunits and differential distribution of expression of catalytic subunits of CK2α and CK2α’ in the human body. A and B were created in BioRender (White, A. (2025) eqznm6m https://BioRender.com/). **(C)** Averaged expression of CSNK2A2 in non-cognitively impaired (NCI) and dementia samples from the Allen Brain Institute: Aging, Dementia and TBI study in the parietal cortex (PCx), Temporal cortex (TCx), and Hippocampus (HIP) are shown (patients with TBI were excluded from the analyses). **(D, E)** Quantification of the expression of CSNK2A2 and CSNK2A1 in PCx from NCI and dementia from the Allen Brain Institute: Aging, Dementia and TBI study (n=20-27/group). Significance determined by unpaired t-test, *p<0.05. **(F)** Immunoblot for CK2α’ and CK2α in protein lysates from frontal cortex of age and sex matched healthy controls (C) and patients with FTD (left panel). Arrow indicates CK2α’ specific band, asterisk indicates unspecific band. (Right panel) Immunoblot for CK2α’ in WT and CK2α’ ^(-/-)^ mice showing antibody-band specificity. **(G, H)** CK2α’ (G) and CK2α (H) protein levels were normalized to GAPDH and shown relative to age/sex matched control (n=7/group). Statistical analyses were conducted using unpaired t-test with welch’s correction p=0.0377. Data are shown as mean <u>±</u> SEM. (**I**) Relationship between CK2α’ levels from (**F**) and the presence of Tau pathology reported by the NIH biobank. Horizontal line represents the group CK2α′ intensity average normalized by GAPDH. (**J**) Immunoblot for CK2α’ and CK2α in the hippocampus of symptomatic WT and PS19 mice. (**K, L**) CK2α’ (K) and CK2α (L) protein levels were analyzed and normalized to GAPDH (n=3 mice/genotype). **(M)** Csnk2a2 in situ hybridization images from symptomatic WT and PS19 hippocampus. Panels show 10x images of hippocampal cell layers with Csnk2a2 RNA probe and DAPI, scale bar=50 μm. (**N, O**) Quantification of Csnk2a2 puncta in the CA1 (p=0.0163) (**N**), and DG (p=0.0003) (**O**) quantified within the cell layer identified by DAPI, (n=3-4 mice/genotype). **(P)** Csnk2a2 in situ hybridization and NeuN immuno-fluorescence images from symptomatic WT and PS19 CA1. Scale bar= 20 μm. **(Q)** Csnk2a2 RNA in situ hybridization and Iba1 immunofluorescence in symptomatic WT and PS19 hippocampus. Top: representative image of single Iba1+ cell in the Stratum Radiatum (CA1). Scale bar is 20 μm. Bottom: 3D rendering of cells Iba1+ cells containing Csnk2a2 puncta showing puncta reside within the microglia cell. **(R)** Quantification of the number of Csnk2a2+ puncta/Iba1+ soma (p<0.0001,n=3-4 mice/group, 8-9 cells/mouse, every data point is a cell). All Data shown are mean <u>±</u> SEM. Statistical analyses were conducted using unpaired t-test and displayed with compact letter display.

To examine whether CK2α’ was also elevated in mouse models of tauopathy we utilized the PS19 mouse model, which expresses human tau with a P301S mutation associated with familial forms of FTD and other tauopathies (*38*). Pathology and neuronal loss in PS19 mice are seen largely in the hippocampus, although it does spread to other brain regions including the neocortex, entorhinal cortex and amygdala (*38, 39*). CK2α’, but not CK2α, was found elevated in the hippocampus of symptomatic (10-12 months) PS19 (*38, 40, 41*) mice by both immunoblotting and RNA in situ hybridization (**Fig. 1J-R**). CK2α’ was elevated in all regions of the hippocampus specially in the CA1 and DG regions (**Fig. 1M-O**), and signal colocalized with NeuN staining, pointing to enhanced expression in neurons (**Fig. 1P**). Enhanced expression of CK2α’ was also observed outside the neuronal layer in microglia (Iba1+) (**Fig. 1Q, R**). Single cell RNA-Seq studies available in The Alzheimer’s Cell Atlas (TACA) (*42–46*) confirmed increased *CSNK2a2* expression in patients with AD in both microglia and neurons in the occipital cortex (OL) and entorhinal cortex (EC) respectively (**Fig. S1**). However, studies in prefrontal cortex (PFC) and superior frontal gyrus (SFG) within the frontal lobe revealed enhanced expression of *CSNK2a2* in cells other than neurons and microglia, suggesting a cell-specific upregulation of *CSNK2a2* that seems to be brain region-dependent. Overall, these results demonstrate a specific upregulation of CK2α’ in brain affected regions of patients with dementia and in hippocampal neurons and microglia of mouse models of tauopathy.

### Genetic depletion of Protein kinase CK2α’ impacts tau pathology *in vitro* and *in vivo*

Previous studies have shown that CK2 holoenzyme regulates tau phosphorylation in various cell and mouse models of AD (*16, 47*), but whether CK2α’ subunit specifically contributes to this phenomenon is unknown. We first explored the impact of silencing CK2α or CK2α’ in Neuro2a (N2a) cells expressing a pathological tau variant (Tau-P301L) in Tau phosphorylation. N2a cells were treated with either scrambled RNA (ScrRNA) or silencing RNA for CK2α (siCK2α) or CK2α’ (siCK2α’) and transfected with either Tau-P301L-EGFP (*48*) or EGFP empty vector (control) (**Fig. 2A**). The AT8 phospho-tau (pTau) antibody was then used to examine the impact of silencing CK2α and CK2α’ on tau pathology (**Fig. 2B**). First, we confirmed the efficacy of the siRNAs decreasing the expression of CK2α or CK2α’ without altering the other subunits (**Fig. 2C-F**) and the increased expression of Tau in N2A transfected cells (**Fig. 2G**). Importantly, siCK2α’ reduced AT8 levels in Tau-P301L transfected cells (p=0.0130) but not siCK2α (p=0.9722) (**Fig. 2B, H**). This data demonstrates an impact of the CK2α’ subunit specifically on pTau accumulation *in vitro*.

**Figure 2.**
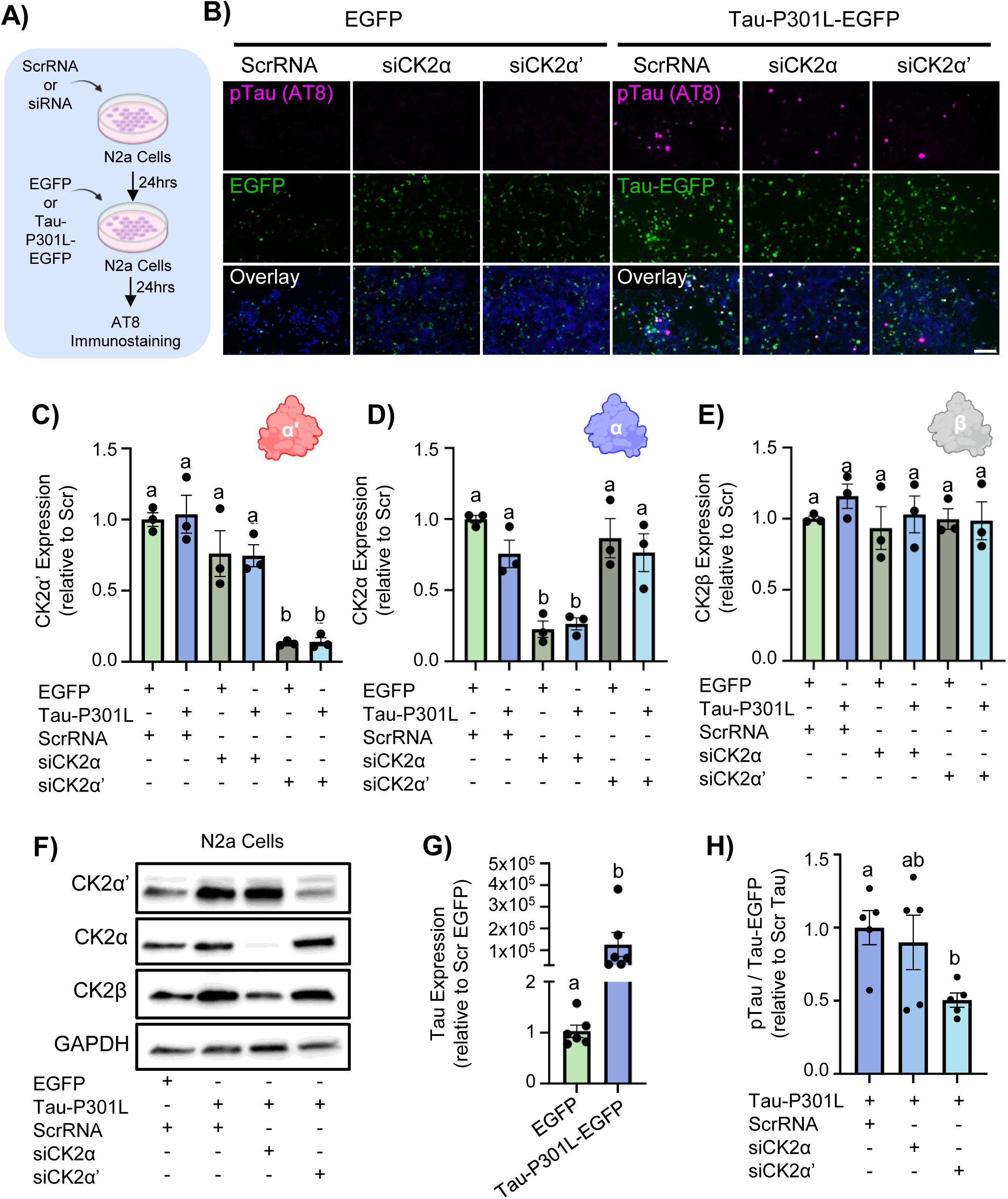
Silencing CK2α’ reduces pTau levels in N2a cells expressing Tau-P301L. **(A)** Schematic of experimental design for N2a cell transfection and immunostaining. **(B)** Representative images of AT8 immunostaining and EGFP fluorescence in N2a cells transfected with either EGFP or Tau-P301L-EGFP and treated with ScrRNA, siCK2α, or siCK2α’. Scale bar= 100 μm. **(C-E)** RT-qPCR analysis of mRNA expression levels of CK2α’ **(C)** (Scr-EGFP vs. siCK2α’-EGFP p=0.0003, Scr-EGFP vs. siCK2α’-Tau-P301L p=0.0003, Scr-Tau-P301L vs. siCK2α’-EGFP p=0.0002, Scr-Tau-P301L vs. siCK2α’-Tau-P301L p=0.0002, siCK2α-EGFP vs. siCK2α’-EGFP p=0.0049, siCK2α-EGFP vs. siCK2α’-Tau-P301L p=0.0056, siCk2α-Tau-P301L vs. siCK2α’-EGFP p=0.0059, siCk2α-Tau-P301L vs. siCK2α’-Tau-P301L p=0.0067), CK2α **(D)** (Scr-EGFP vs. siCK2α-EGFP p=0.0008, Scr-EGFP vs. siCk2α-Tau-P301L p=0.0013, Scr-Tau-P301L vs. siCK2α-EGFP p=0.0161, Scr Tau-P301L vs. siCk2α Tau-P301L p=0.0262, siCK2α EGFP vs. siCK2α’ EGFP p=0.004, siCK2α-EGFP vs. siCK2α’-Tau-P301L p=0.0144, siCk2α-Tau-P301L vs. siCK2α’-EGFP p=0.0065, siCk2α-Tau-P301L vs. siCK2α’-Tau-P301L p=0.0235), and CK2β **(E)** in transfected N2a cells. Expression levels were normalized to GAPDH and presented relative to ScrRNA-EGFP condition (n=3 independent replicas). **(F)** Immunoblot of CK2α’, CK2α and CK2β in N2A cells transfected with EGFP or Tau-P301L-EGFP and treated with siCK2α’ or siCK2α. (**G**) RT-qPCR analysis of mRNA expression levels of huTau expression in N2a cells transfected with EGFP or Tau-P301L-EGFP transfected N2a cells. Expression levels were normalized to GAPDH and presented relative to EGFP condition (p=0.0464, n=6/condition). **(H)** Quantification of AT8+ cells normalized to the number of EGFP+ cells, presented relative to ScrRNA-Tau-P301L condition. (n=4/condition; Scr-Tau-P301L vs. siCK2α’-Tau-P301L p=0.013). All data shown as mean ± SEM. Statistical analyses were conducted using one-way ANOVA with tukey’s multiple and displayed with compact letter display.

We then generated a PS19 mouse haploinsufficient for CK2α’ (PS19;CK2α’^(+/-)^) (**Fig. S2A-C**) to assess the impact of depleting CK2α’ on tau pathology *in vivo* in both prodromal (∼7 months) and at a late symptomatic age (∼10-12 months) (*38*) (**Fig. 3A-B**). In the prodromal age no significant differences were observed in the accumulation of AT8 in PS19 and PS19;CK2α’^(+/-)^ in all three examined regions of the hippocampus (CA1, CA3, and DG) and overlaying cortex (**Fig. 3C-E, Fig. S2D-F**). As expected, in the late symptomatic groups we observed a significant increase in AT8+ area in both the PS19 and PS19;CK2α’^(+/-)^ mice in the CA1, CA3, and DG (**Fig. 3C-E**) and cortex (**Fig. S2D-F)** relative to all other groups. However, PS19;CK2α’^(+/-)^ mice showed a significant decrease in AT8 signal compared to PS19 in the CA1, DG, and cortex (CA1; p=0.0075, DG; p=0.0186 and Cortex; p<0.0001) (**Fig. 3C,E, Fig. S2D-F**) suggesting an overall decrease in pTau upon depletion of CK2α’. Similar results were obtained using the AT100 phospho-Tau antibody (**Fig. S2G-J**). Decreased pTau in PS19;CK2α’^(+/-)^ was also confirmed by immunoblotting from whole hippocampal samples at 9 months (**Fig. 3F-I**).

**Figure 3.**
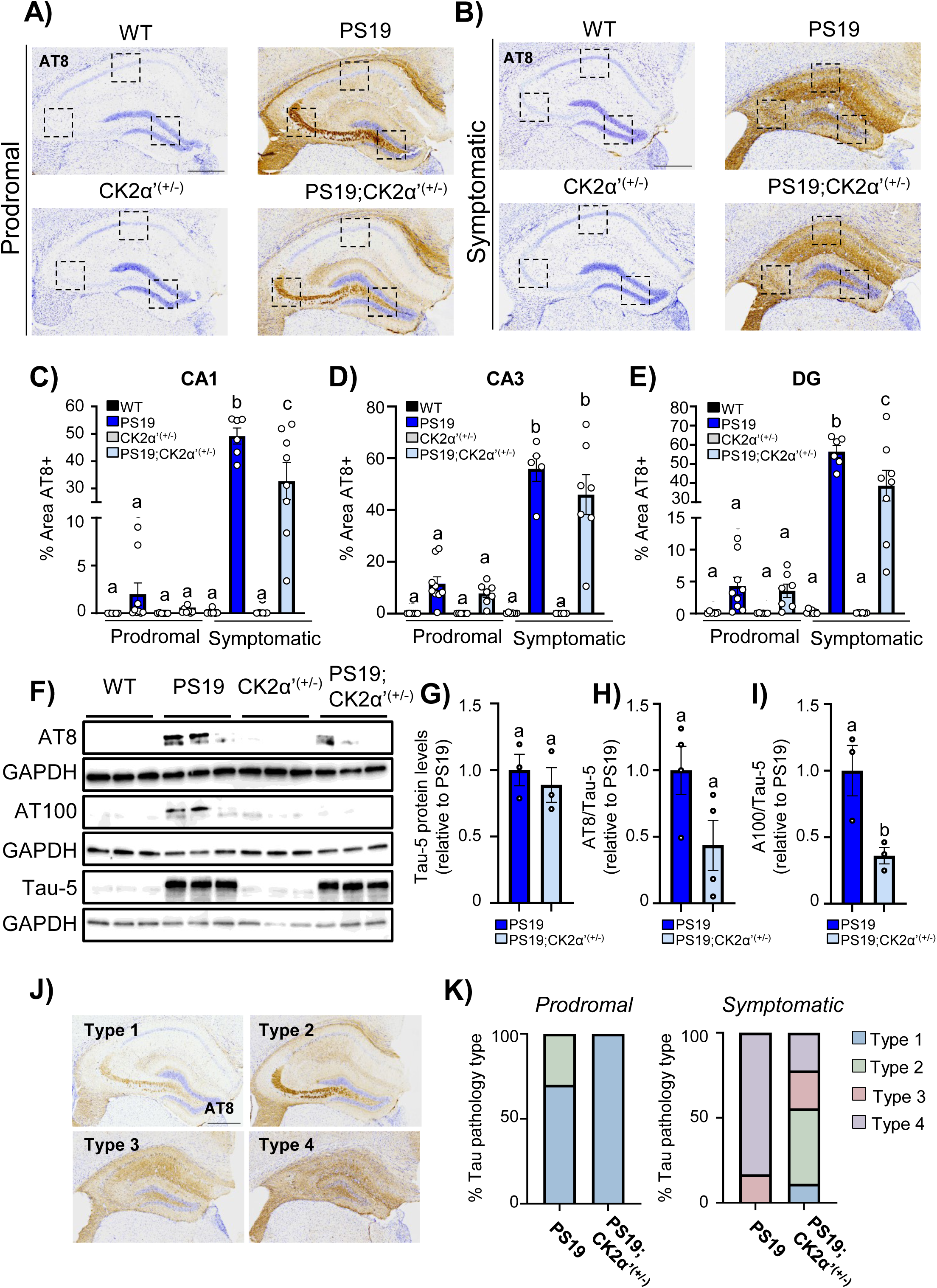
CK2α’ haploinsufficiency reduced tau pathology in PS19 mice. **(A-B)** Representative hippocampal sections immunostained for pTau (AT8;Ser202/Thr205). Scale bar=500 μm. **(C-E)** Quantification of AT8+ immunoreactivity (% area) in the CA1 (C), CA3 (D) and DG (E) regions in prodromal and symptomatic cohorts (n=6-9 mice/genotype). Data represented as mean <u>±</u> SEM. Statistical analyses were conducted using two-way ANOVA with tukey’s posthoc analysis and displayed with compact letter display. (**CA1 significant comparisons**: prodromal: WT vs. symptomatic: PS19 p<0.0001, prodromal: WT vs. symptomatic: PS19;CK2α’^(+/-)^ p<0.0001, prodromal: PS19 vs. symptomatic: PS19 p<0.0001, prodromal: PS19 vs. symptomatic: PS19;CK2α’^(+/-)^ p<0.0001, prodromal: CK2α’^(+/-)^ vs. symptomatic: PS19 p<0.0001, prodromal: CK2α’^(+/-)^ vs. symptomatic: PS19;CK2α’^(+/-)^ p<0.0001, prodromal: PS19;CK2α’^(+/-)^ vs. symptomatic: PS19 p<0.0001, prodromal: PS19;CK2α’^(+/-)^ vs. symptomatic: PS19;CK2α’^(+/-)^ p<0.0001, symptomatic: WT vs. symptomatic: PS19 p<0.0001, symptomatic: WT vs. symptomatic: PS19;CK2α’^(+/-)^ p<0.0001, symptomatic: PS19 vs. symptomatic: CK2α’^(+/-)^ p<0.0001, symptomatic: PS19 vs. symptomatic: PS19;CK2α’^(+/-)^ p=0.0075, symptomatic: CK2α’^(+/-)^ vs. symptomatic: PS19;CK2α’^(+/-)^ p<0.0001; **CA3 significant comparisons:** all significant comparisons p<0.0001; **DG significant comparisons:** prodromal: WT vs. symptomatic: PS19 p<0.0001, prodromal: WT vs. symptomatic: PS19;CK2α’^(+/-^p<0.0001, prodromal: PS19 vs. symptomatic: PS19 p=<0.0001, prodromal: PS19 vs. symptomatic: PS19;CK2α’^(+/-)^ p<0.0001, prodromal: CK2α’^(+/-)^ vs. symptomatic: PS19 p<0.0001, prodromal: CK2α’^(+/-)^ vs. symptomatic:PS19;CK2α’^(+/-)^ p<0.0001, prodromal: PS19;CK2α’^(+/-)^ vs. symptomatic: PS19 p<0.0001, prodromal :PS19 CK2α’^(+/-)^ vs. symptomatic: PS19;CK2α’^(+/-)^ p<0.0001, symptomatic: WT vs. symptomatic: PS19 p<0.0001, symptomatic: WT vs. symptomatic: PS19;CK2α’^(+/-)^ p<0.0001, symptomatic: PS19 vs. symptomatic: CK2α’^(+/-)^ p<0.0001, symptomatic: PS19 vs. symptomatic :PS19;CK2α’^(+/-)^ p=0.0186, symptomatic: CK2α’^(+/-)^ vs. symptomatic:PS19;CK2α’^(+/-)^ p<0.0001). **(F)** Immunoblot for AT8 (pS202/T205), AT100 (pT212/pS214), and Tau-5 (Total Tau) in hippocampal samples obtained from 9 month mice. **(G-I)** Quantification of Tau-5 (G), AT8/Tau-5 (H) and AT100/Tau-5 (I) from images shown in F (n=3-4 mice/genotype). All data are shown as mean <u>±</u> SEM. **(J)** Representative pTau pathology types. **(K)** Proportion of each pTau pathology type in prodromal and symptomatic cohorts (n=7-10 mice/genotype). Fisher’s exact t-test 7m p=0.11 & 12m p=0.0587.

Tau pathology has been previously reported to follow a characteristic progressive pattern of deposition in the hippocampus that can be categorized into 4 pathology types (*49–52*) (**Fig. 3J**). These pathology types have been shown to correlate with hippocampal volume (*50, 52*) and offer a more holistic pathological assessment. We observed differential distributions of patterns between PS19 and PS19;CK2α’^(+/-)^ mice with a more dramatic alteration in late symptomatic stages where nearly all PS19 mice presented a type 4 pathology in contrast to the PS19;CK2α’^(+/-)^ that presented a variation in types with almost 50% of mice presenting type 2 pathology (prodromal p=0.11, and symptomatic p=0.0587) (**Fig. 3K**). The observed modulation of tau pathology underscored a critical role for CK2α’ in the modulation of tau phosphorylation and associated pathology.

### RNA-Seq analyses revealed CK2**α**’ alters the expression of genes related to immune response and synaptic function

To determine whether the reduction in Tau pathology translates into wider changes in cellular signaling and gene regulation, we next analyzed RNA-seq profiles from CK2α’-deficient mice (**Fig. 4**). We focused on the prodromal phase to identify pathways involved in disease onset that may be missed if only later stages of the disease are studied where significant neuronal loss is present. Gene expression analyses in the PS19 vs WT mice identified 477 significant DGEs (**Fig. 4A**). Gene ontology pathway analysis revealed that many of the DGEs belonging to the top eight dysregulated pathways corresponded to immune and inflammatory functions (**Fig. 4B**), as previously reported (*53, 54*). Other important GO pathways were associated with synaptic dysfunction where the top 5 corresponded to dysregulation in synapse assembly, regulation of synapse organization, regulation of synapse structure or activity, post synapse organization and synaptic pruning (**Fig. 4B**). Synaptic pruning was upregulated in PS19 vs WT, whereas the other synaptic pathways related to organization and regulation of activity were largely downregulated (**Fig. 4C**).

**Figure 4.**
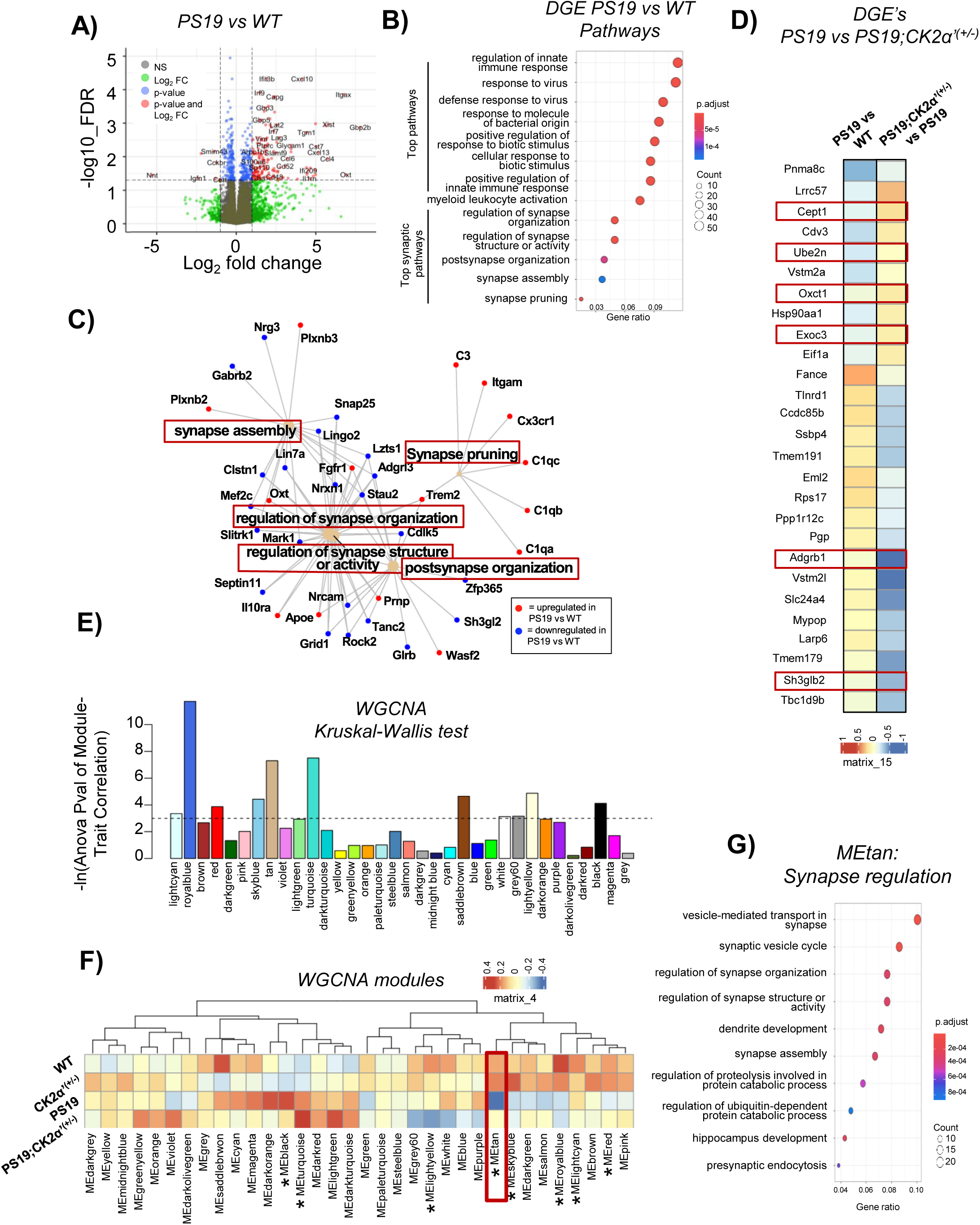
CK2α’ haploinsufficiency reduced transcriptional alterations in apoptotic/phagocytic and synaptic-function related genes in PS19 mice. **(A)** Volcano plot of DGEs in PS19 vs WT comparison (n=3-4 mice/genotype). **(B)** Gene-ontology showing the top biological pathways for DGEs in PS19 vs WT. **(C)** Gene network for top 5 synaptic pathways impacted in PS19 vs WT. Red dots indicate genes upregulated in PS19 mice. Blue dots indicate genes down-regulated in PS19 mice. **(D)** Heat map displaying DGEs identified between PS19 and PS19;CK2α’^(+/-)^ mice (n=3-4 mice/genotype). Red boxes indicate genes with apoptotic/phagocytic and immune-related functions. *q>0.05. **(E)** WGCNA identified modules, dotted line indicates significance as determined by the Kruskal wallis test between WT, CK2α’^(+/-)^, PS19, and PS19;CK2α’^(+/-)^ mice (n=3-5 mice/genotype). **(F)** Heatmap of modules detected from WGCNA indicating differences across genotypes. * indicates p<0.05 determined by Kruskal-wallis test. **(G)** Gene ontology for top biological pathways for MEtan module.

Direct comparison of PS19 vs PS19;CK2α’^(+/-)^ mice yield a small set of 26 significant DGEs (**Fig. 4D**). Several of these DGEs were related to Apoptotic/phagocytic and Immune functions (*Cept1, Ube2n, Oxct1, Exoc3, Adgrb1, Sh3glb2*). Most of the DGEs between PS19 and PS19;CK2α’^(+/-)^ were up-regulated in PS19 mice compared to WT but reduced expression in PS19;CK2α’^(+/-)^. One example Adgrdb1, an adhesion-GPCR, strongly associated with the engulfment and pruning of synapses (*55, 56*), was significantly downregulated in PS19;CK2α’^(+/-)^ mice (**Fig. 4D**). We then assessed whether the manipulation of CK2α’ had a broader effect on modules of genes with similar biological functions despite not reaching the strict >Log2 criteria. Weighted Gene Co-Expression Network Analysis (WGNCA) revealed 34 gene modules with 8 modules (red, lightcyan, royalblue, skyblue, tan, lightyellow, turquoise, and black) showing statistical significance detected via a Kruskal-Wallis test among our groups (p<0.05) (**Fig. 4E, File S2**). Among the 8 significant gene modules the red, tan, and black modules showed an expression pattern in the PS19;CK2α’^(+/-)^ mice that more closely resembled the expression pattern of WT mice showing an overall rescue in expression compared to PS19 mice (**Fig. 4F**). We focused our analyses on the tan module for being the most significant gene module among the three rescued modules (**Fig. 4E**). The tan module composed on 215 genes (Kruskal-wallis p=0.016) (**File S2**) was represented by pathways associated with synaptic function and assembly, with top pathways being vesicle-mediated transport in synapse, synaptic vesicle cycle, and regulation of synapse organization (**Fig. 4G**). The overall expression of genes associated with these pathways presented a depletion in the PS19 mice that was ameliorated in the PS19;CK2α’^(+/-)^ group (**Fig. 4F**). Overall, these results suggested an amelioration in synapse pruning and an improvement in synaptic function and organization when deleting CK2α’ in PS19 mice.

### PS19 mice lacking CK2**α**’ ameliorated microglia reactivity, neuroinflammation and phagocytic activity

pTau is a potent driver of microglia reactivity (*9, 38, 57–60*) which in turn contributes to exacerbate Tau-mediated pathology (*53, 61–65*). Having established that CK2α’ is elevated in PS19 microglia, and that deletion CK2α’ decreased phospho-Tau accumulation and resulted in the downregulation of genes associated with apoptosis/phagocytosis and synaptic pruning, we investigated whether these phenotypes were associated with changes in microglia reactivity. We first examined the number of Iba1+ microglia in the hippocampus of mice as an indicator of microgliosis (**Fig. 5A, B, Fig. S3A-E**). Iba1+ cells significantly increased in PS19 mice compared to WT, especially in symptomatic mice. Importantly, PS19;CK2α’^(+/-)^ significantly reduced the number of Iba1+ microglia in the CA1 and CA3 (**Fig. 5A, B, Fig. S3A-E**). Microgliosis was also reduced in the cortex of PS19;CK2α’^(+/-)^ mice compared to PS19 (**Fig. S3F-H**). We also analyzed the number of GFAP+ cells in the hippocampus of PS19 mice in both age groups (**Fig. S4**), as increased astrogliosis is also reported in PS19 mice (*38*). However, no significant differences were found between PS19 and PS19;CK2α’^(+/-)^ mice (**Fig. S4**) suggesting that CK2α’ mediated changes in glial cells are specific to microglia.

**Figure 5.**
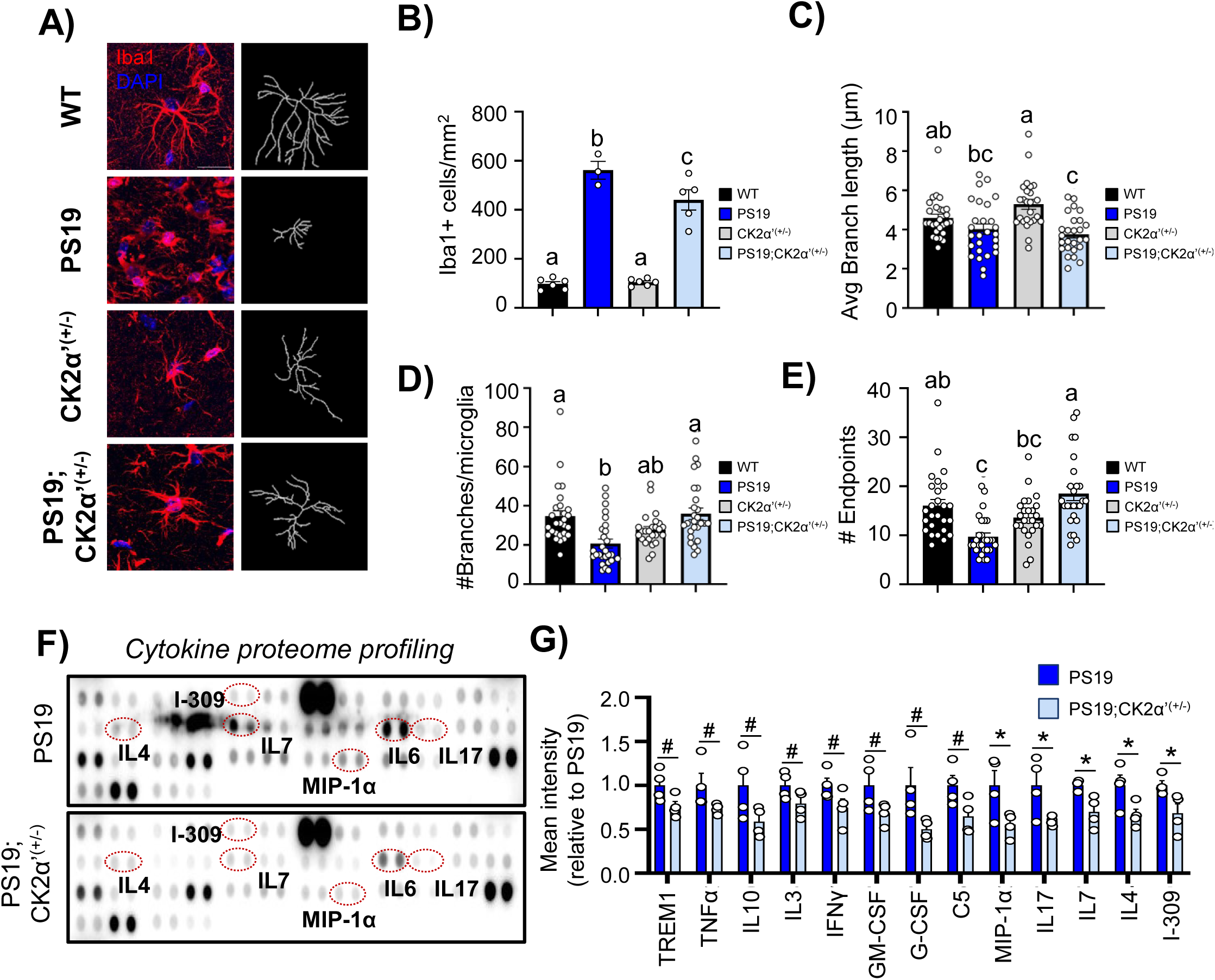
CK2α’ haploinsufficiency impacts Iba1+ microglia morphology and cytokine production in PS19 mice. **(A)** Iba1+ microglia immunostaining and microglia skeleton representation from the CA1 of symptomatic animals used for microglia morphology analysis. **(B)** Quantification of the number of Iba1+ positive cells in the CA1 (n=3-6 mice/genotype; WT vs. PS19 p<0.0001, WT vs. CK2α’^(+/-)^ p=0.9963, WT vs. PS19; CK2α’^(+/-)^ p<0.0001, PS19 vs. CK2α’^(+/-)^ p<0.0001, PS19 vs. PS19;CK2α’^(+/-)^ p=0.0313, CK2α’^(+/-)^ vs. PS19;CK2α’^(+/-)^ p<0.0001). **(C)** Quantification of microglia branch length (WT vs. PS19 p=0.4402, WT vs. CK2α’^(+/-)^ p=0.1371, WT vs. PS19; CK2α’^(+/-)^ p=0.041, PS19 vs. CK2α’^(+/-)^ p=0.0184, PS19 vs. PS19;CK2α’^(+/-)^ p=0.927, CK2α’^(+/-)^ vs. PS19;CK2α’^(+/-)^ p=0.0004). **(D)** Number of branches per microglia (WT vs. PS19 p=0.0008, WT vs. CK2α’^(+/-)^ p=0.2116, WT vs. PS19;CK2α’^(+/-)^ p=0.9908, PS19 vs. CK2α’^(+/-)^ p=0.3063, PS19 vs. PS19;CK2α’^(+/-)^ p=0.0114, CK2α’^(+/-)^ vs. PS19; CK2α’^(+/-)^ p=0.0757). **(E)** Number of endpoints per microglia (WT vs. PS19 p=0.0003, WT vs. CK2α’^(+/-)^ p=0.3541, WT vs. PS19;CK2α’^(+/-)^ p=0.5111, PS19 vs. CK2α’^(+/-)^ p=0.0556, PS19 vs. PS19;CK2α’^(+/-)^ p=0.0004, CK2α’^(+/-)^ vs. PS19;CK2α’^(+/-)^ p=0.0179) (n=27 cells (3 mice/genotype, 3 slices/mouse, 3 cells/slice). **(F)** Representative cytokine arrays from PS19 and PS19;CK2α’^(+/-)^ at 9 months from hippocampal protein extracts. Cytokines that showed significant differences are circled in red. **(G)** Selected quantifications of cytokines showing significance or trending significance. Quantifications are represented relative to WT (n=4 mice/genotype;Trem1 p=0.0624, Tnfα p=0.1009, IL10 p=0.0948, IL3 p=0.0952, IFNy p=0.0972, GM-CSF p=0.0816, G-CSF p=0.0562, C5 p=0.0522, Mip 1α p=0.0482, IL17 p=0.0386, IL7 p=0.0481, IL4 p=0.046, I309 p=0.039). Data are shown as mean <u>±</u> SEM. Statistical analyses were conducted using ANOVA with Geisser-greenhouse correction and Tukey’s post-hoc analysis in **B-E** and unpaired t-test in **G.** Significance displayed with compact letter display or #p≤0.1, *p<0.05.

We then performed morphological assessments of microglia using measurements corresponding to ramification and correlating with microglia state (**Fig. 5A, C-E**). It has been previously reported that microglia adopt an amoeboid morphology in PS19 mice, characterized by the loss of microglia processes, and that is associated with a more reactive and phagocytic state (*9, 66*). Both the PS19 and PS19;CK2α’^(+/-)^ mice showed a decrease in the average branch length of microglia in the CA1 (**Fig. 5C**). However, PS19;CK2α’^(+/-)^ mice showed a significant increase in the number of branches and number of endpoints compared to the PS19 mice (**Fig. 5D,E**) indicating an improvement in the morphological state of microglia.

Since the morphological state of microglia is strongly related to its reactive state and neuroinflammation we first assessed the expression of a variety of cytokines and other inflammatory molecules associated with microglia reactivity by using a cytokine array panel with hippocampal extracts from symptomatic PS19 and PS19;CK2α’^(+/-)^ (**Fig. 5F, G**). Importantly, we confirmed that several cytokines were significantly decreased (p<0.05) in PS19;CK2α’^(+/-)^ mice relative to PS19 including, MIP-1α, IL17, IL7, IL4, and I309 with several others showing trending decreases (0.05<p<0.1) including TREM1 and TNFα (**Fig. 5F, G**). We also investigated whether changes in overall cytokine levels and microglia reactivity upon CK2α’ depletion related to an altered phagocytic state. For this we assessed the levels of CD68, a lysosomal/endosomal marker strongly associated with phagocytic function of microglia and normalized CD68 levels to the number of Iba1+ cells observed in each animal and time point (**Fig. 6A-E, Fig. S3I**). We observed increased levels of CD68 in all regions of the hippocampus in the PS19 mice at both prodromal and symptomatic stages (**Fig. 6C-E**). PS19;CK2α’^(+/-)^ mice also presented increased CD68 levels compared to WT but displayed significantly less CD68 in the CA1 and a trending decrease in the CA3 and DG at the prodromal stage compared to PS19 mice (**Fig. 6C-E**). Importantly, symptomatic PS19;CK2α’^(+/-)^ mice displayed significantly less CD68 in all hippocampal regions compared to PS19 mice (**Fig. 6C-E**). Expression of another phagocytic marker Clec7a was also significantly reduced in PS19;CK2α’^(+/-)^ compared to PS19 mice supporting the impact of CK2α’ deletion in the amelioration of microglia phagocytosis (**Fig. 6F-H**).

**Figure 6.**
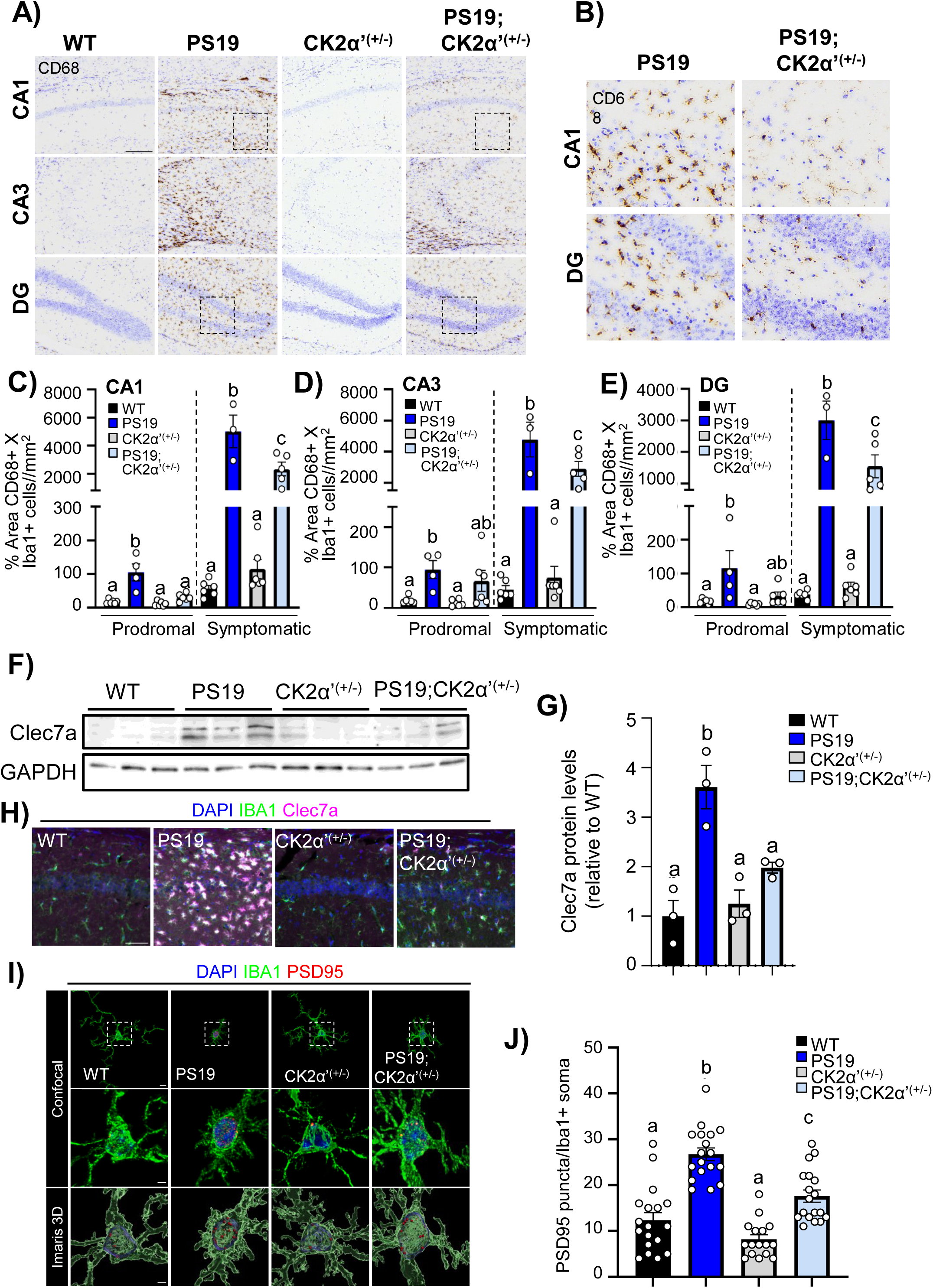
CK2α’ haploinsufficiency decreases microglia reactivity and synaptic engulfment. **(A)** CD68 immunostaining with cresyl violet counterstain in the hippocampus of symptomatic mice. Scale bar=150 μm. (**B**) zoomed images from A in the CA1 and DG. **(C-E)** Quantifications of % area CD68+ normalized by Iba1+ cell count in prodromal and symptomatic cohorts in CA1, CA3 and DG (n=3-6 mice/genotype). Comparisons shown relative to cohort age group (**CA1:** *Prodromal:* WT vs. PS19 p<0.0001, PS19 vs. CK2α’^(+/-)^ p<0.0001, PS19 vs. PS19;CK2α’^(+/-)^ p=0.0003, *Symptomatic:* WT vs. PS19 p<0.0001, WT vs. PS19;CK2α’^(+/-)^ p=0.0036, PS19 vs. CK2α’^(+/-)^ p<0.0001, PS19 vs. PS19;CK2α’^(+/-)^ p=0.0042, CK2a’(+/-) vs. PS19;CK2α’^(+/-)^ p=0.0045 **CA3:** *Prodromal:* WT vs. PS19 p=0.0328, WT vs. PS19;CK2α’^(+/-)^ p=0.1739, PS19 vs. CK2α’^(+/-)^ p=0.0193, PS19 vs. PS19; CK2α’^(+/-)^ p=0.6895, CK2α’(+/-) vs. PS19;CK2α’^(+/-)^ p=0.1045 *Symptomatic:* WT vs. PS19 p<0.0001, WT vs. PS19;CK2α’^(+/-)^ p=0.0003, PS19 vs. CK2α’^(+/-)^ p<0.0001, PS19 vs. PS19;CK2α’^(+/-)^ p=0.0404, CK2α’^(+/-)^ vs. PS19; CK2α’^(+/-)^ p=0.0003 **DG:** *Prodromal:* WT vs. PS19 p=0.0176, WT vs. PS19; CK2α’^(+/-)^ p=0.9281, PS19 vs. CK2α’^(+/-)^ p=0.0091, PS19 vs. PS19;CK2α’^(+/-)^ p=0.0536, CK2α’^(+/-)^ vs. PS19;CK2α’^(+/-)^ p=0.7762 *Symptomatic:* WT vs. PS19 p<0.0001, WT vs. PS19;CK2α’^(+/-)^ p=0.0018, PS19 vs. CK2α’^(+/-)^ p<0.0001, PS19 vs. PS19;CK2α’^(+/-)^ p=0.011, CK2α’^(+/-)^ vs. PS19; CK2α’^(+/-)^ p=0.002. **(F, G)** Immunoblotting and quantification of protein levels for Clec7a in the hippocampus of symptomatic animals (n=3 mice/genotype). (**H**) Representative immunofluorescence of Clec7a in the hippocampus of symptomatic animals. **(I)** Iba1 and PSD95 immunostaining from the CA1 of symptomatic animals. Top row: whole microglia including processes, scale bar= 5μm. Middle row: higher magnification view of the soma (white dotted box in top row), scale bar= 2μm. Bottom row: 3D reconstruction of Iba1 and PSD-95 signal generated from Imaris Software, scale bar= 2μm. **(J)** Quantification of the number of PSD-95 puncta/Iba1+ soma (n= 16-18 cell, 3 mice/genotype, 2-3 slices/mouse, 22 cells/slice; WT vs. PS19 p<0.0001, WT vs. PS19; CK2α’^(+/-)^ p=0.0854, PS19 vs. CK2α’^(+/-)^ p<0.0001, PS19 vs. PS19; CK2α’^(+/-)^ p=0.0025, CK2α’^(+/-)^ vs. PS19;CK2α’^(+/-)^ p=0.0208). Data are show as mean ± SEM. Statistical analyses were conducted using one-way ANOVA with Tukey’s post-hoc displayed with compact letter display.

Aberrant microglia-mediated synaptic pruning has been extensively reported in PS19 mice and contributes to synaptic dysfunction and neuronal loss (*53, 61–65*). Considering the changes observed in CD68 and Clec7a markers of microglia phagocytosis and transcriptional changes related to apoptosis and synaptic pruning upon deletion of CK2α’ we assessed whether PS19;CK2α’^(+/-)^ microglia decreased synaptic engulfment. To quantify synapse phagocytosis/engulfment we co-stained for the postsynaptic marker PSD-95 and Iba1 (*67, 68*) and examined the CA1 stratum radiatum of symptomatic mice (**Fig. 6I**). PS19 mice exhibited a significant increase in PSD-95+ puncta within Iba1+ cells relative to control mice (**Fig. 6I, J**), demonstrating increased phagocytosis in PS19 microglia. Importantly, PS19;CK2α’^(+/-)^ showed a significant reduction in PSD-95 engulfment (**Fig. 6I, J**) indicating an amelioration in the phagocytic activity of microglia upon deletion of CK2α’.

### CK2**α**’ happloinsufficiency improved hippocampal synaptic density and synaptic function in PS19 mice

We then assessed whether the observed changes in synaptic gene expression and synaptic engulfment by microglia upon deletion of CK2α’ were translated into enhanced synaptic density and neuronal activity in PS19 mice. We first conducted NeuN immunostaining analyses in the hippocampus of PS19 and PS19;CK2α’^(+/-)^ mice to determine if depletion of CK2α’ could impact the overall neuronal loss characteristic of PS19 mice (*38, 69–71*). Significant depletion of NeuN+ cells and hippocampal atrophy was only seen in PS19 groups at a symptomatic age (**Fig. 7A, B, Fig. S5A-C**). Importantly, PS19;CK2α’^(+/-)^ mice revealed a significant increase in the number of NeuN+ cells compared to PS19 in the CA1 (**Fig. 7A, B, Fig. S5A-C**).

**Figure 7.**
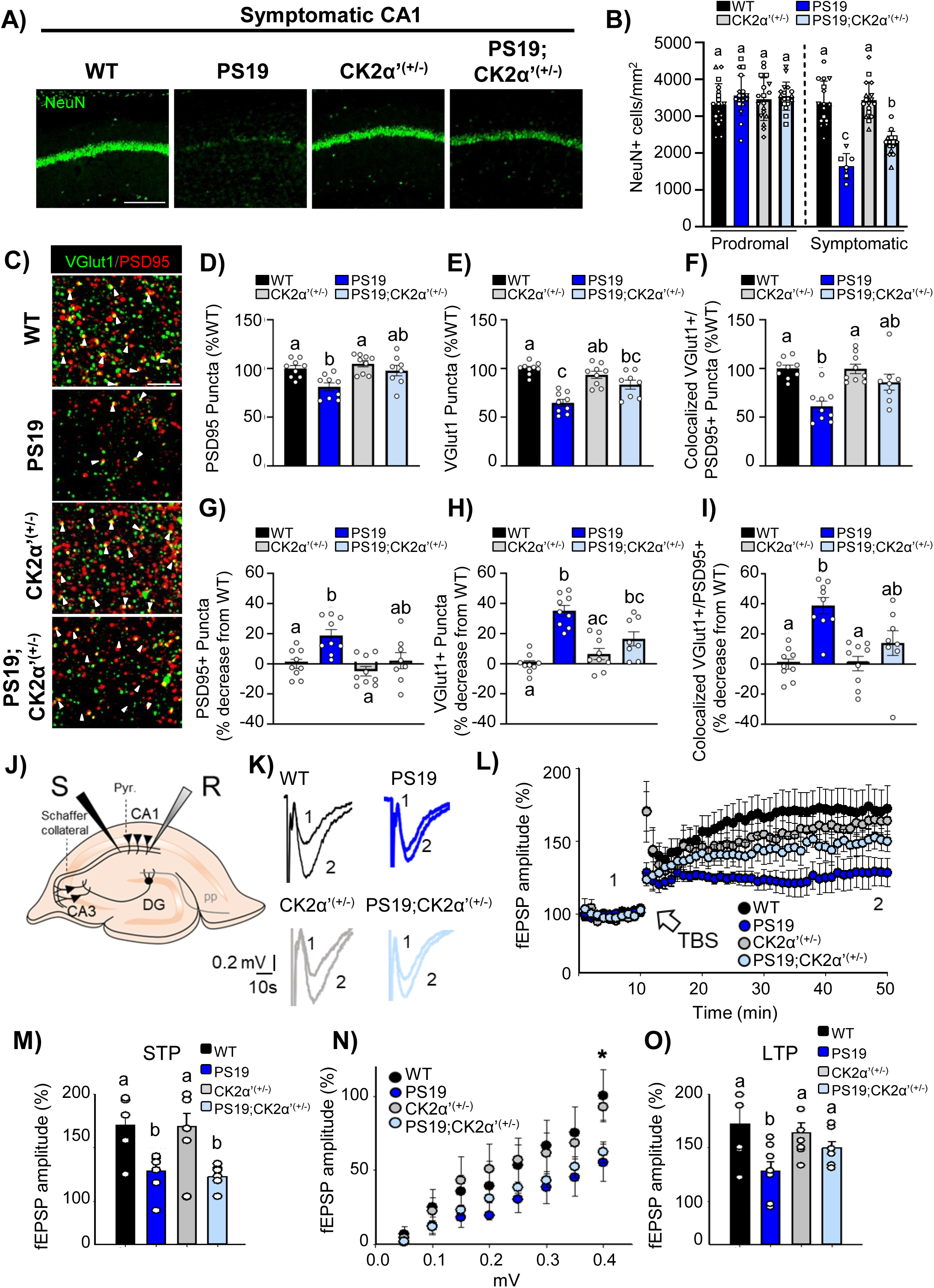
CK2α’ haploinsufficiency partially rescued hippocampal synapse loss and improved LTP PS19 mice. **(A)** Representative NeuN staining in the CA1 of symptomatic mice. Scale bar=200 μm. **(B)** Quantification of NeuN+ cells (cells/mm^2^) in the CA1 of prodromal and symptomatic cohorts (n= 3-6 mice/genotype; 2-3 slices/mouse; 1 data point= 1 slice, slices from same animal share a shape). Comparisons shown relative to age group (Significant comparisons: *Symptomatic:* WT vs. PS19 p<0.0001, WT vs. PS19;CK2α’^(+/-)^ p=0.0002, PS19 vs. CK2α’^(+/-)^ p<0.0001, CK2α’^(+/-)^ vs. PS19;CK2α’^(+/-)^ p=0.0001). **(C)** Vglut1 and PSD95 immunostaining in the CA1 of symptomatic mice. White arrow heads indicate colocalization. Scale bar= 2.5 μm. **(D-F)** Quantification of PSD95 puncta (WT vs. PS19 p=0.0445, PS19 vs. CK2α’^(+/-)^ p=0.0008) (**D**), Vglut1 puncta (WT vs. PS19 p=0.0003, WT vs. PS19;CK2α’^(+/-)^ p=0.0212, PS19 vs. CK2α’^(+/-)^ p<0.0001) (**E**) and Vglut1/PSD95 colocalized puncta (WT vs. PS19 p=0.0017, PS19 vs. CK2α’^(+/-)^ p=0.0004) (**F**) relative to WT. (n=3 mice/genotype (3 slices/mouse, 3 images/slice, 1 data point=slice average). **(G-I)** Analyses from (D-F) represented as a % decrease from WT (G: WT vs. PS19 p=0.0445, PS19 vs. CK2α’^(+/-)^ p=0.0008 H: WT vs. PS19 p=0.0003, WT vs. PS19;CK2α’^(+/-)^ p=0.0212, PS19 vs. CK2α’^(+/-)^ p<0.0001, I: WT vs. PS19 p=0.0017, PS19 vs. CK2α’^(+/-)^ p=0.0004) **(J)** Experimental set-up: R, recording electrode; S stimulating electrode. **(K)** Representative traces before (1) and after (2) Theta-burst stimulation (TBS). **(L)** Course temporal of TBS application inducing LTP in pyramidal neurons of CA1 (n=6-7 mice/genotype). **(M)** Short-term potentiation (STP) results, significance reported relative to WT (WT vs. PS19 p=0.014, WT vs. PS19;CK2α’^(+/-)^ p=0.007). **(N)** Input-output curves. Averaged fEPSP amplitude, (n=3mice/genotype). **(O)** Long-term potentiation (LTP) results, significance reported relative to WT (WT vs. PS19 p=0.007). All data are shown as mean <u>±</u> SEM. Statistics: one way-ANOVA with Tukey’s (**B**), repeated measures ANOVA with Geisser-greenhouse correction and Tukey’s (**D-I**), or one-way ANOVA with Holm-Sidak (**M**-**O**). Significance displayed as compact letter display.

We then evaluated if total synapse density was also positively altered in the CA1. We conducted immunostaining and colocalization analyses for the vesicular glutamate transporter 1 (VGlut1) (pre-synaptic marker of excitatory synapses), and the postsynaptic marker PSD-95 (**Fig. 7C-I**). As expected, PS19 mice showed a significant decrease in the number of synapses (colocalized puncta VGlut1-PSD95) compared to WT mice (**Fig. 7F, I**). The reduction in colocalized puncta in PS19 mice was due to a concomitant reduction in both PSD-95 and VGlut1 markers (**Fig. 7D, E, G, H**). In PS19;CK2α’^(+/-)^ mice we observed a decrease in the % loss of both PSD-95 (**Fig. 7D, G**) and VGlut1 (**Fig. 7E, H**). This resulted in the amelioration of total synapse loss in PS19;CK2α’^(+/-)^ mice and obtained values for colocalization were no longer significant respect to WT (**Fig. 7F, I**). These results demonstrate a partial rescue in the density of excitatory synapses in the CA1 of PS19;CK2α’^(+/-)^ mice aligning with the amelioration of hippocampal atrophy observed in this region (**Fig. 7B**) and the reduced synapse engulfment mediated by microglia (**Fig. 6I, J**). In line with these results, silencing CK2α’ in primary cortical neuron cultures transfected with Tau-P301L-EGFP increased spine density compared to scramble (**Fig. S6**). To investigate functional implications of the rescue seen in PS19;CK2α’^(+/-)^ hippocampal atrophy and synaptic density we assessed hippocampal synaptic plasticity by performing extracellular field excitatory postsynaptic potential (fEPSP) recordings in the CA1 region of acute hippocampal slices (**Fig. 7J, K**). Theta-burst stimulation (TBS) at the Schaffer collaterals induced short-term potentiation (STP), which resulted in a significant increase in synaptic efficacy at CA1 synapses in both WT and CK2α’^(+/-)^ mice, as evidenced by the enhanced fEPSP amplitude (170 ±13%, n = 6; 170 ± 12%, n = 7) (**Fig. 7L, M**). In contrast, the increase in fEPSP amplitude was significantly reduced in PS19 and PS19;CK2α’^(+/-)^ mice relative to WT controls post TBS (123 ± 4%, n = 7; 128 ± 7%,n = 6; **Fig. 7L, M**), indicating impaired STP in these models. Furthermore, we examined long-term potentiation (LTP) at CA3-CA1 synapses, a form of synaptic plasticity linked to learning and memory. It has been previously shown that PS19 mice present impaired LTP in the CA1-CA3 pathway (*38, 72, 73*). First, we observed significant differences in input-output curves at CA3-CA1 synapses between PS19 and PS19;CK2α’^(+/-)^ mice when compared to WT, which suggests that excitatory synaptic input properties were altered in these models (**Fig. 7N**). TBS at the Schaffer collaterals induced robust LTP in both WT (173 ± 12%, n = 6) and CK2α’^(+/-)^ (164 ± 9%, n = 7; **Fig. 7L, O**). However, in the PS19 mouse model, the magnitude of LTP was significantly diminished compared to WT (128 ± 9%, n = 7; vs. WT p=0.007) (**Fig. 7L, O**), suggesting a disruption in the molecular mechanisms underlying LTP. Importantly, in the PS19;CK2α’^(+/-)^ mice, LTP was partially restored (150 ± 6%, n = 6; vs. WT p=0.199) (**Fig. 7L, O**), pointing to a potential compensatory mechanism that partially mitigates the LTP deficits observed in the PS19 mice. Importantly, paired-pulse facilitation (PPF) was unaltered across all experimental groups, suggesting that the observed changes in synaptic plasticity were most likely due to postsynaptic mechanisms dependent on the hippocampus (**Fig. S7**).

### CK2**α**’ haploinsufficiency improved PS19 behavior on Barnes Maze

Finally, we assessed whether the overall amelioration in Tau-pathology, microgliosis and synaptic dysfunction mediated by CK2α’ had an impact in cognitive behaviors. Mice were trained on the Barnes maze task for 5 consecutive days on the location of a target hole (training) and on the 6^th^ day a probe trial was conducted with the target hole removed (**Fig. 8A**). During the training trials the primary latency or the time to first locate the escape hole was recorded (**Fig. 8B-C**). All mice in the prodromal group demonstrated an improvement on time needed to locate the escape hole from day 1 vs day 5 (WT p=0.0272; PS19 p=0.0.0067; CK2α’^(+/-)^ p=0.0002; PS19;CK2α’^(+/-)^ p<0.0001) with no significant difference between the genotypes (**Fig. 8B**). All mice in the symptomatic group also showed a capability of acquiring the task demonstrating a significant improvement from day 1 vs day 5 (WT p<0.0001; PS19 p=0.0109; CK2α’^(+/-)^ p<0.0001; PS19;CK2α’^(+/-)^ p=0.0037). However, PS19 and PS19;CK2α’^(+/-)^ showed a significant impairment compared to WT, starting on day 3 for PS19 (p=0.0009) and day 4 for PS19;CK2α’^(+/-)^ (p=0.0306) (**Fig. 8C**). The training data suggested that both symptomatic PS19 and PS19;CK2α’^(+/-)^ mice showed similar impaired learning on the Barnes Maze compared to WT mice, although deficits in PS19;CK2α’^(+/-)^ were attenuated.

**Figure 8.**
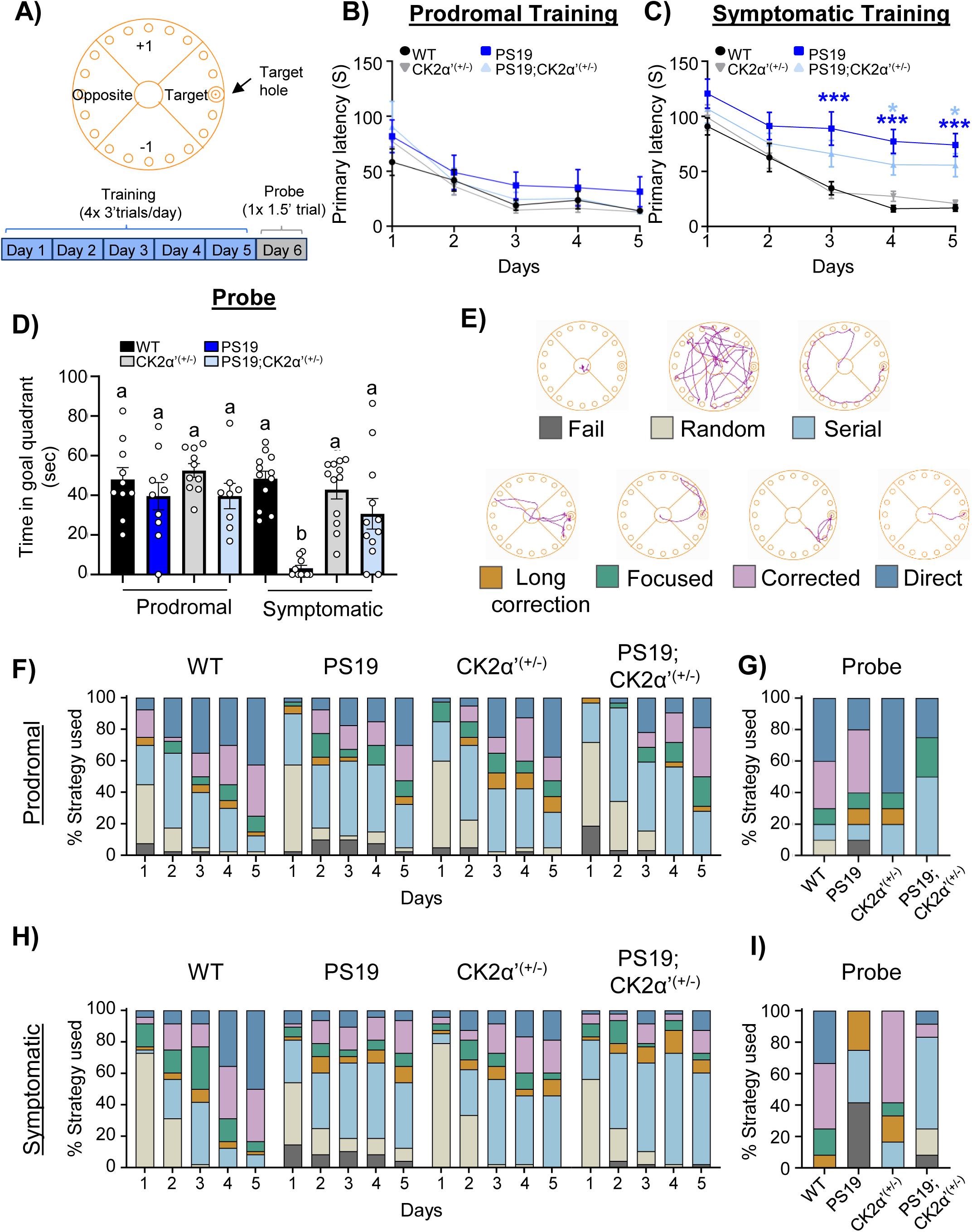
Depletion of CK2α’ improved spatial memory of PS19 mice on the Barnes maze. **(A)** Schematic of the Barnes Maze divided into 4 quadrants (Target, Opposite, +1 and -1) and experimental set up. **(B-C)** Primary latency (s), or time to first explore escape hole at prodromal time point. Statistical analyses were conducted using two-way ANOVA with Bonferroni’s multiple comparison’s. Prodromal; genotype p=0.0573, training day p<0.0001 and interaction p=0.9690. Symptomatic: genotype p<0.0001, training day p<0.0001 and interaction p=0.7018. Differences for each training day were conducted using Bonferroni’s multiple comparisons and * show significance relative to WT. **(D)** Time spent in goal quadrant of Barnes maze during probe trial at prodromal and symptomatic time points. n=10-12 mice/genotype, 2 outliers were removed from PS19 group using ROUT (Q=1%). **(E)** Representative search strategy traces based on their navigation patterns on the maze. **(F,H)** Spatial strategies used to locate the escape hole across training days for each genotype at prodromal and symptomatic stages, respectively. **G,I)** Spatial strategies used to locate the previously learned escape target hole during probe trial. Data are shown as mean ± SEM. Statistical analyses were conducted using Two-way ANOVA with Bonferroni’s multiple comparisons (B-C) or Two-Way ANOVA with Tukey’s post-hoc analysis (D). Significance displayed with either *p<0.05, **p<0.01, ***p<0.001, ****p<0.0001 or compact letter display.

Following the training phase of the Barnes maze we conducted a probe phase in which the escape hole was covered and the time spent exploring the escape hole quadrant was recorded. At the prodromal age there was no significant difference in time spent in the escape hole quadrant for any genotype (**Fig. 8D**). At the symptomatic age, the PS19 mice showed a significant impairment in the time spent in the goal quadrant compared to WT (p<0.0001) (**Fig. 8D**). Importantly, the PS19;CK2α’^(+/-)^ mice spent significantly more time than the PS19 mice in the goal quadrant with no significant difference from WT (vs. PS19 p=0.0113 vs. WT p=0.2335) demonstrating a rescue in spatial memory for the PS19;CK2α’^(+/-)^ on the probe phase of the Barnes maze (**Fig. 8D**).

The PS19 mouse model is known to present motor deficits due to hindlimb paralysis and weakness (*38*), which could confound Barnes Maze results if PS19 mice perform more poorly due to less distance traveled. Prodromal mice displayed no difference in distance traveled on Barnes Maze (**Fig. S8A**). While the symptomatic PS19 mice traveled less distance in the Barnes Maze compared to WT mice, they did not show a significant difference in distance traveled compared to PS19;CK2α’^(+/-)^ mice (**Fig. S8B**). In addition, in open-field testing we found no differences between groups at the prodromal time point (**Fig. S8C**). On the contrary, symptomatic mice showed significant hyperactivity in the open field task, demonstrating the ability of PS19 mice to move (**Fig. S8D**). Indeed, no significant differences were observed in the open field between PS19 and PS19;CK2α’^(+/-)^ mice (**Fig. S8D**). This data supports the improvement in cognitive abilities seen in the PS19;CK2α’^(+/-)^ mice are not due to changes in motor function but rather are related to an improvement in memory.

Mice can use different cognitive strategies to navigate the Barnes Maze and locate the escape hole (*74*). As the mouse learns the task they show improvement in the strategy used to locate the escape hole (*74*) (**Fig. 8E**). Assessing these cognitive strategies is important to provide a deeper insight into how memory and spatial learning are being utilized. All animals in the prodromal group demonstrated a similar pattern in spatial strategies over the training phase (**Fig. 8F**) and showed a predominance in the corrected and direct strategies in the probe phase (**Fig. 8G**). Symptomatic PS19 and PS19;CK2α’^(+/-)^ mice both showed impaired spatial strategies over the training phase, by day 5 both genotypes largely displayed either failing, random or serial strategies (**Fig. 8H**). However, during the shortened probe phase, PS19 displayed a worsened overall strategy, with predominant failing, serial, and long correction strategies compared to PS19;CK2α’^(+/-)^ mice that displayed more serial, corrected and direct strategies (**Fig. 8I**). Overall, the PS19;CK2α’^(+/-)^ mice displayed improvement in the probe phase of the Barnes Maze in both the time spent in the escape quadrant and strategy used to locate the target, indicating improvements in spatial memory.

## Discussion

Tauopathies are a heterogeneous group of neurodegenerative disorders for which effective treatments are still not available, in part due to an incomplete understanding of the underlying mechanisms. In this study, we present a novel target, CK2α’, a catalytic subunit of CK2, as a potential upstream regulator of tau mediated neurodegeneration, acting at the intersection of neuroimmune signaling and synaptic dysfunction.

Here, we report that CK2α’ expression is abnormally elevated in the brains of dementia patients and in the hippocampus of PS19 mice, with preferential upregulation observed in hippocampal neurons and microglia. These findings are supported by multiple single-cell RNA-sequencing (snRNA-seq) studies in human AD, which consistently show increased CK2α’ expression in excitatory neurons (*44–46*). Additionally, the study by Gerrits et al. (*43*) in the occipital lobe (OL) has reported preferential CK2α’ expression in AD-associated microglia. Other scRNA-seq studies have detected elevated CK2α’ expression in various brain cell types, with patterns varying by dataset and brain region (*44–46*). On the other hand, CK2α seems to be increased in cortical astrocytes of the 5x FAD and PS19 models (*75*). This collective evidence suggests that CK2α’ dysregulation in AD/ADRD may be both cell-type- and region-specific. Consistent with our findings, a concurrently submitted study has reported that CK2α′ is elevated in AD models and contributes to APP-regulated glial inflammation (*76*), independently supporting and strengthening our conclusions.

Increased CK2 levels have been associated with tau pathology (*16, 17, 47, 77*), microglia state (*78, 79*) and general inflammation (*80–86*). However, many of these studies were conducted in cell models (*16, 17, 34, 47*) and genetic evidence for the specific contributions of the individual CK2 subunits to tau pathology remained largely unexplored. Our studies *in vitro* using N2A cells transfected with Tau-P301L showed that silencing CK2α’, but not CK2α, decreased pTau accumulation. Furthermore, our studies *in vivo* reducing the levels of CK2α’ in the PS19 model also influenced tau pathology by decreasing the accumulation of pTau and tau burden in the hippocampus and cortex of PS19;CK2α’^(+/-)^ mice. In support to our data, previous studies *in vitro* showed that overexpression of CK2α’ subunit decreased the activity of I2PP2A/SET, a cellular protein that functions as an endogenous inhibitor of protein phosphatase 2A (PP2A), increasing pTau. In contrast overexpression of CK2α increased the activity of I2PPA2a/SET (*77*). Although early studies demonstrated that Tau is a physiological substrate of CK2, at least in specific contexts such as neurogenesis (*87*), the findings by Perez et al. (*77*) suggest that the effects we observed after manipulating CK2α’ on pTau accumulation could be mediated through intermediary molecules that modulate Tau phosphorylation.

Haploinsufficiency of CK2α’ also increased the expression of synaptic genes, synaptic density, and synaptic function in the PS19 mice. Along these lines, haploinsufficiency of CK2α’ in a mouse model of HD also showed improved synaptic density and function (*15, 88*), although how exactly CK2α’ influences these processes is still unknown. A previous study showed that CK2α’ can influence the expression of synaptic genes, especially those related to glutamatergic excitatory signaling (*15*). A more recent study showed that pharmacological inhibition of CK2 mitigated AD tau pathology by preventing the NMDA receptor subunit NR2B synaptic mislocalization (*47*). NR2B mediates long-term depression (LTD), and LTD specifically induces the phosphorylation of tau. These studies support our findings and strengthen the relationship between CK2α’, tau phosphorylation, and the regulation of neuronal function in both AD and other neurodegenerative diseases. However, since CK2α’ is found elevated in both neurons and microglia, it is still unknown whether the effects mediated by CK2α’ in pTau accumulation and synaptic function are cell or non-cell autonomous, warranting further investigation.

CK2 has been identified as a mediator of microglia reactivity and inflammation in different models (*78, 79*) and it has been well-established as a mediator of inflammation in a variety of contexts including SARS-CoV infection, cancers, bacterial infections, intestinal inflammation and renal failure (*80–85*). This is largely mediated via CK2’s involvement in regulating several large inflammatory factors such as NF-κB, STAT1, and EGR-1 (*86*). It is known that chronic activation of NF-kB increases the accumulation of pTau (*9*). On the other hand, pathological tau has also been shown to induce inflammation and NF-kB activation by interacting directly with inflammatory receptors such as toll-like receptors 2 (TLR2) (*10, 11*) creating a vicious cycle of worsening inflammation and tau pathology (*9–12*). Our data demonstrated that CK2α’ haploinsufficiency significantly reduced the number of Iba1+ cells in the hippocampus, reduced the levels of the microglia reactive and phagocytic markers CD68 and Clec7a, ameliorated microglia morphological changes associated with pathology, reduced the levels of pro-inflammatory cytokines in PS19 mice and reduced microglia-mediated synaptic engulfment. In support to our findings, previous studies in human microglia derived from hiPSCs treated with CK2 inhibitors also reduced cytokines production (*79*). Therefore, all together, these data support a specific key role of CK2α’ in mediating neuroinflammation via activation of microglia.

Loss of synaptic density and neuronal dysfunction in AD models has been previously connected to increased phagocytic microglia (*39, 63, 89, 90*). Our results demonstrated a decrease in microglia phagocytosis of the PSD-95 synaptic marker in PS19;CK2α’^(+/–)^ mice suggesting that CK2α’ is influencing the phagocytosis of synapses by microglia, but the mechanisms behind this are yet to be elucidated. CK2α’ could facilitate these processes by regulating the transcription of genes involved in phagocytosis and synaptic engulfment and/or by phosphorylating and activating specific proteins related to these processes. Deletion of CK2α’ in PS19 mice resulted in the down-regulation of several genes related to apoptosis/phagocytosis and synapse engulfment such as *Cept1, Ube2n, Oxct1, Exoc3, Adgrb1* and *Sh3glb2* (*55, 56, 91–95*) which can contribute to the phenotypes observed in PS19;CK2α’^(+/–)^ mice. It is also possible that CK2α’ phosphorylates elements of the complement component such as C9 and C1r, known CK2 substrates (*96, 97*) and influence microglia-mediated synaptic pruning.

In conclusion, our findings identified CK2α’ as a central and previously underappreciated regulator of tauopathy pathogenesis, acting at the crossroads of neuroimmune signaling, synaptic function, and tau phosphorylation. By demonstrating that CK2α’ haploinsufficiency alleviates tau pathology, reduces neuroinflammation, and improves synaptic integrity and function in PS19 mice, we provide compelling genetic evidence that CK2α’ contributes to multiple pathological hallmarks of tau-driven neurodegeneration. Importantly, these effects appear to occur through complex, context-dependent mechanisms that may differ across cell types and disease stages. The observation that CK2α’ influences both neuronal and microglial compartments underscore the need to dissect its cell-autonomous versus non-cell-autonomous roles. Given the growing interest in CK2 as a therapeutic target and the limitations of current inhibitors that lack subunit specificity, the development of CK2α’-selective modulators (*76, 98*) may represent a promising avenue for disease-modifying therapies in tauopathies and related neurodegenerative conditions. Future studies elucidating the molecular and cellular pathways downstream of CK2α’ will be critical for translating these findings into clinical applications.

## Materials and Methods

### Study design

In this study we report the upregulation of CK2α’ in AD human samples and mouse models and investigated the role of CK2α’ in tauopathy. We hypothesized that CK2α’ contributes to tau-associated pathology and that reducing its levels would improve outcomes in tauopathy models. Data was collected from AD/ADRD post-mortem human tissue and AD mouse and cell models. Methods used in this study include western blotting, immunostaining (tissue and cells), bulk RNA-Seq, electrophysiology and behavioral assays. Publicly available human genomic datasets were also analyzed, as described.

For mouse experiments, animals were assigned to cohorts at birth based on genotype and aged to predetermined time points; no additional randomization was performed. Sample sizes for all experiments were chosen based on previous literature and our prior experience with similar experiments; no formal power calculations were performed, exact sample sizes are detailed in figure legends. Equal numbers of male and female mice were used. Data is presented as mouse averages unless otherwise detailed in the figure legend.

Experimenters were not blinded during behavioral testing, however all image analysis was performed on blinded samples to reduce experimenter bias.

### Cell culture: Culturing, transfections, and immunocytochemistry

Neuro-2a cells (N2a; ATCC CCL-131) were cultured in DMEM with 10% FBS and penicillin/streptomycin at 37 °C and 5% CO_₂_. For silencing experiments, cells were plated in 12-well plates or Matrigel-coated coverslips. Cells were transfected with siRNA targeting CK2α or CK2α′ (Qiagen) or a negative control using DharmaFECT 1 (25 nM). After 24 h, cells were transfected with pEGFP-C1 or Tau-P301L-EGFP using jetOptimus and processed 24 h later. For immunocytochemistry, cells were fixed, blocked in 5% NGS/TBST, incubated with AT8 primary antibody overnight, followed by Alexa-fluor secondary antibodies, and mounted with ProLong Gold for imaging (see Supplementary Methods for primary neuron culturing details;(*99*)).

### Mouse Lines

For this study we used the B6;C3-Tg(Prnp-MPAT-P301S)PS19Vle/J (or PS19) (*38*) mouse line (Jackson #008169 maintained with purchased B6C3H breeders; obtained from Dr. Karen Ashe at University of Minnesota), and CK2α’^(+/-)^ mice, originally obtained from Dr. Seldin at Boston University (Taconic biosciences TF3062) (*31*) and maintained on the C57Bl/6J background for several generations. We crossed PS19 and CK2α’^(+/-)^ mice and this cross generated 4 genotypes of interest WT (PS19^0/0^;CK2α’^(+/+)^), CK2α’^(+/-)^ (PS19^0/0^;CK2α’^(+/-)^), PS19 (PS19^Tg/0^;CK2α’^(+/+)^), and PS19;CK2α’^(+/-)^ (PS19^Tg/0^;CK2α’^(+/-)^) mice. For all experiments littermate WT and CK2α’^(+/-)^ controls were used. Two time points were used in experiments; pre-symptomatic (6-7 months) and symptomatic (10-12 months). All mice were housed under standard SPF conditions. All animal care and sacrifice procedures were approved by the University of Minnesota Institutional Animal Care and Use Committee (IACUC) in compliance with the National Institutes of Health guidelines for the care and use of laboratory animals under the approved animal protocol 2307-A41243.

### Human postmortem tissue

Postmortem human frontal cortex (Brodmann’s area 46) FTD patients and controls subjects was provided by NeuroBioBank of National Institutes of Health. A total of 6 female (3 control/3FTD) and 8 male (4 control/4 FTD) samples were used, and samples were age and sex matched. Ages ranged from 60-77 with the average age of all samples being 70. More detailed information on specific samples can be found in **file S1**.

### Immunoblotting

Human and mouse protein was extracted as previously described (*14*). Tissue was homogenized in 25□mM Tris-HCl pH 7.4, 150□mM NaCl, 1□mM EDTA, 0.1% SDS, 1% Triton X-100, incubated on ice 15-30 min, then SDS was added to 2% and samples were heated at 100°C for 5 min. Lysates were centrifuged at 4°C, 13,000 g for 20 min and supernatants collected. Protein concentration determined by BCA assay (Pierce).

Protein samples were separated on 4–20% SDS Criterion TGX Stain-Free gels (BioRad) and transferred to a 0.2□μm nitrocellulose membrane in Tris–Glycine Buffer (25□nM Tris-Base, 200□mM Glycine) at 25□V for 30□min (Trans Turbo Transfer system). Membranes were blocked in 5% milk in TBST (0.5% Tween-20) for 1□hour, incubated overnight at 4□°C in primary antibody diluted in 2.5% milk in TBST, then incubated in HRP-conjugated secondary antibody (Amersham 1:5000) for 1□hour. Bands were visualized with SuperSignal Chemiluminiscent substrate (Thermo Scientific) using a GE ImageQuant Las4000 imager. Primary antibodies: CK2α’ (1:2000; Rabbit, Novus NB100-379), CK2α (1:1000; Rabbit, Abcam ab76040), Clec7a (1:500; Rat invivoGen mabg-mdect), At8 (1:1000; Mouse, Invitrogen MN1020), At100 (1:1000; Mouse, Invitrogen MN1060), Tau-5 (1:1000; Mouse, Abcam ab80579), and GAPDH (1:5000; Mouse, Santa cruz sc-365062).(See supplementary Methods for cytokine immunoblotting details)

### Immunohistochemistry

Immunohistochemistry experiments were conducted as previously described (*14, 100, 101*) using fluorescent or 3,3’-diaminobenzidine (DAB) detection. Mice were anesthetized (Avertin, 250 mg/kg) and perfused intracardially with TBS (25 mM Tris-base, 135 mM Nacl, 3 mM KCl, pH 7.6) supplemented with 7.5 mM heparin. Brains were fixed with 4% PFA at 4°C for 4–5 days, cryoprotected with 30% sucrose for 4–5 days and embedded in a 2:1 mixture of 30% sucrose in TBS:OCT (Tissue-Tek). Coronal sections (16 μm) were stored in a 50% glycerol-50% at -20°C. Three hippocampal sections were analyzed per mouse.

For immunofluorescent staining, sections were blocked in 5% normal goat serum (NGS) in TBST for 1 h at room temperature. Primary antibodies were incubated overnight at 4°C in TBST containing 5% NGS. Secondary Alexa-fluorophore-conjugated antibodies (Invitrogen) were added (1:200 in TBST with 5% NGS) for 1 h at room temperature. Slides were mounted in ProLong Gold Antifade with DAPI (Invitrogen) and subsequently imaged. Primary antibodies used and dilutions are as follows: GFAP (1:2000; Chicken Millipore Sigma AB5541), Iba1 (1:500; Rabbit Fujifilm Wako 019-19741), Iba1 (1:500, Goat Fujifilm Wako 011-27991), NeuN (1:1000; Mouse Millipore MAB377), Clec7a (1:1000;Rat inviogen mabg-mdect), PSD95 (1:500; Rabbit Invitrogen 51-6900) and Vglut1 (1:500; Guinea pig Millipore AB5905). Imaging of immunofluorescent slices was conducted using a confocal microscope (Stellaris, Lecia). For NeuN, Iba1, and GFAP imaging 20x tiles were acquired with a 16um z-dimension and 1um step size. For PSD95 and VGlut1 images were acquired as described for synapse analysis.

For DAB staining antigen retrieval was performed using either Tris-EDTA pH=9.0 (Biolegend 422703) or Rodent decloaker (Biocare medical RD913) at 80°C for 30 minutes. Sections were blocked at room temperature in 10% NGS/TBST for 1 hour, then incubated in primary antibodies overnight at 4C in 5%NGS/TBST. Sections were incubated in biotin-conjugated secondaries (Jackson Immuno Research Labs Biotin-SP (long spacer) AffiniPure) in 5%NGS/TBST for 1 hour, followed by 3% hydrogen peroxide for 20 minutes. Sections were incubated in tertiary antibodies (VECTASTAIN Elite ABC, HRPS kit, Vector laboratories) in 5%NGS/TBST for 1 hour per manufacturer instructions. DAB chromogen diluted in DAB substrate buffer (Biolegend) was applied for sufficient staining. Sections were counterstained with 0.1% cresly violet, dehydrated, cleared in ethanol/xylene and mounted in Permount mounting medium (Fisher scientific). Primary antibodies used and dilutions are as follows: pTau (Ser202, Thr205) AT8 (1:1000; Mouse, Invitrogen Mn1020), pTau (Ser214, Thr212) AT100 (1:5000; Mouse Invitrogen MN1060) and CD68 [RM1031] (1:2500; Rabbit, Abcam ab303565). DAB images were captured at 10x (Echo Revolve) and at 40x (EasyScane One, Motic)(See Supplementary Methods for Tau pathology type assessment;(*49–52*)). All analyses were conducted in either Qupath (*102, 103*) or ImageJ (FIJI) (*104*) (See Supplementary Methods for Image Analysis details, including microglia morphology analysis (*105, 106*))

### In situ hybridization

RNAscope was conducted using a custom Csnk2a2 probe (ACDbio protocol). Slides were then blocked and incubated overnight with anti-IBA1. The next day, Alexa-fluorophore secondary antibodies were applied, and slides were mounted with ProLong Gold Antifade containing DAPI for imaging.(See Supplementary Methods for details).

### Synapse analysis

Synapse analysis was conducted as previously described (*14, 88, 101, 107, 108*). Immunohistochemical staining was conducted as described above with Vglut1 and PSD95, with blocking increased to 20% NGS+TBST for 2 hours and primary and secondary antibodies diluted in 10% NGS+TBST. Fluorescent images from the molecular layer of the CA1 were taken on a confocal microscope (Stellaris, Lecia) at 63x with z-step dimension of 0.34 μm with 15 steps were generated. Lecia stellaris software lightning processing was applied to images post-acquisition and edited images were used for analysis. Maximum projections of three sections per slice were generated. Puncta analyses were conducted blinded using the PunctaAnalyzer Plugin (Durham,NC,USA) in FIJI as previously described (*14, 88, 101, 107, 108*). Data is represented as slice averages with three slices per animal.

### Electrophysiology

Mice were anesthetized with isoflurane (2%) and decapitated for slice preparation. Brains were rapidly removed into (sucrose 189 mM, glucose 10 mM, NaHCO3 26 mM, KCl 3 mM, MgSO4 5 mM, CaCl2 0.1 mM, NaH2PO4 1.25 mM). Transverse hippocampal slices (350 μm) were cut on a vibratome (LEICA VT1000S) and incubated at room temperature in oxygenated artificial cerebrospinal fluid (aCSF; NaCl 124 mM, KCl 5 mM, NaH2PO4 1.25 mM, MgSO4 2 mM, NaHCO3 26 mM, CaCl2 2 mM, glucose 10 mM; gassed with 95% O2, 5% CO2; pH=7.4) for 45 min-1 h. Experiments were performed at 30–34°C with continuous perfusion.

Field excitatory postsynaptic potentials (fEPSPs) were recorded in the CA1 region of the hippocampus. fEPSPs in were evoked with a stimulating electrode placed on the Schaffer collateral (0.33 Hz) using brief current pulses (200 μs, 0.1–0.2 mA). Extracellular recording electrodes were filled with aCSF. Stimulation was adjusted to obtain a fEPSP peak amplitude of approximately 1 mV during control conditions. After a stable fEPSP baseline period of 10 min, LTP was induced by a Theta-burst stimulation (TBS) consisting of a series of 10 bursts of 5 stimuli (100 Hz within the burst, 200 ms interburst interval), which was repeated 4 times (5 s apart). Data were filtered at 3 kHz and acquired at 10 kHz using pCLAMP 10.2 software (Molecular Devices, RRID: SCR_011323).

A stimulus-response curve (0.05-0.4 mV, mean of five fEPSPs at each stimulation strength) was compiled for the different mice used. For paired-pulse ratio experiments, two fEPSPs were evoked 40 ms apart for 0.5 min at baseline frequency (6 times) at the beginning of the baseline recording. The PPR was expressed as the amplitude of the second fEPSP divided by the amplitude of the first fEPSP.

### Behavior

Behavioral testing was performed with the support from the Universtiy of Minnesota Mouse Behavior Core. Mice were handled daily for ≥ 1 week and habituated to testing room ≥ 30 min before tasks. All tasks were recorded using ANYmaze software (Stoelting Co., Wood Dale, Illinois) and overhead cameras.

Barnes maze was conducted on a 20 hole Barnes Maze (San Diego Instruments) under 450-500lux illumination with black and white visual cues around the room, as described (*74, 109, 110*). Mice were placed in the center under a cover with lights off, lights were turned on and the cover removed within 30 seconds. Training consisted of four 3-minute trials per mouse for 5 days, with 20-30 minutes intertrial intervals. The maze was cleaned with 70% ethanol between mice and rotated between trials. Primary latency, or the time to first explore the escape hole, was recorded. Trials ended when mice entered the escape box or after 3 minutes, at which point the mouse was gently guided to the escape box. During the probe trial, the escape hole was covered and time spent in each maze zone was recorded. Spatial strategy was determined by a blinded investigator.

Open field was conducted in 40cmx40cm boxes under 150-200lux illumination. Mice were placed in boxes and behavior was recorded for 1 hour. Boxes were thoroughly cleaned between mice with 70% ethanol.

### RNA-Seq Analysis

RNA-Sequencing was conducted by Novogene (Sacramento, CA). Gene expression analysis was carried out using the CHURP pipeline (https://doi-org.ezp2.lib.umn.edu/10.1145/3332186.3333156) using n=4-7 mice/genotype with female(F)/male(M) ratios as follows; 4 WT (2F/2M), 5 CK2α’^(+/-)^ (2F/3M), PS19 (3F/4M) (3 high pathology and 4 low pathology), and 4 PS19;CK2α’^(+/-)^ (3F/1M). Differential gene expression was determined with DESeq2 using default setting (*111*) (v1.46.0). Genes with a FDR <=□0.05 were considered significant. Outliers’ identification was performed using Cook’s distance (DESeq2). Driver factors of gene expression variance (genotype and/or sex) were evaluated using R (v4.4.2) package variancePartition (1.36.3). Pathway and clustering analysis were completed gProfiler2 (*112*) (v0.2.3) and clusterProfiler (v4.14.6). Data visualization was done using various R graphic packages, including ggplot2, ggraph, and DESeq2 visualization functions. The RNA-seq data set generated in this manuscript has been deposited at GEO (accession number GSE298505). The reviewer token to access the GEO deposited data is **mlwxmuactlibryl**.(See Supplementary methods for RNA extraction and qPCR details).

### Statistical analyses and data representation

For electrophysiology experiments data were analyzed using Clampfit 10.2 software (pCLAMP, Molecular Devices, RRID: SCR_011323). Data are presented as mean ± SEM. To estimate changes in synaptic efficacy, STP was quantified by comparing the mean fEPSP amplitude during 1 minute after the application of the TBS. LTP was quantified as for STP but 40 minutes after the protocol. Graphs were obtained using SigmaPlot 14.0. Before applying any statistical comparison, the data were subjected to Shapiro-Wilk normality and equal variance tests. For any comparisons between two groups, two-paired Student’s t-test was used. For multiple comparisons to the same control, One-way ANOVA and Holm-Sidak test was used. P-values less than 0.05 were considered statistically significant.

For all other experiments GraphPad Prism (GraphPad, San Diego, CA, USA) was used to create graphs and conduct statical testing. For all other experiments data is represented as mean ± SEM. Statistical analyses were performed using paired Student’s t-test, unpaired student’s t-test, one-way or two-way ANOVA with post-hoc analysis, as indicated in each figure legend. A p-value <0.05 was considered significant.

## Supporting information

Supplementary File S1. Human FTD samples from NBB

Supplementary File S2. RNA-seq Modules

## List of supplemental materials

Supplemental materials and methods

Fig. S1 to S8

Data File S1-S2

## Authors contributions

RGP conceived and directed the project, helped with data analysis and interpretation, figures preparation and manuscript writing. AW and RT conducted in vitro experiments using N2A cells. AW conducted immunoblotting with human samples, generated all mice used in the study, conducted immunostaining for pTau, CD68, and cytokine analyses, performed all behavioral studies, and contributed to preparing all figures and writing the manuscript. PK helped with behavioral data analysis. PG conducted RNA in situ hybridization for CK2a’ and co-staining with NeuN and Iba1, performed microglia morphology analyses, synapse engulfment by microglia, and GFAP and Iba1 staining. PGU assisted with Clec7a immunoblotting. NBR, PG and AF conducted immunostaining for PSD95/VGlut1 and synapse density analyses. SV and RS helped validate siCK2a’ in neurons. JM helped with execution of DAB immunostaining. MKL contributed to experimental supervision and data interpretation. AW and YZ conducted RNAseq experiments and data analyses. RFM conducted electrophysiological studies in the CA1 and AA helped with experimental supervision and data interpretation of such data.

## Data and materials availability

RNA-Seq data generated in this study is available in the Gene Expression Omnibus repository found at https://www.ncbi.nlm.nih.gov/geo/ accession number GSE298505. The reviewer token to access the GEO deposited data is **mlwxmuactlibryl**.

Allan Brain Atlas Aging, Dementia and TBI dataset that supported conclusions of this article is publicly available at https://aging.brain-map.org/overview/home (*37*).

Human brain tissue was received from the NIH NeuroBioBank repository and is available publicly for request at https://neurobiobank.nih.gov/.

scRNA-Seq study data from human samples that contributed to conclusions of this article can be found in The Alzheimer’s Cell Atlas (TACA) https://taca.lerner.ccf.org/ (*42–46*).

## References

1. B. Kovacech, R. Skrabana, M. Novak, Transition of tau protein from disordered to misordered in Alzheimer’s disease. Neurodegener Dis 7, 24–27 (2010).

2. V. M. Lee, M. Goedert, J. Q. Trojanowski, Neurodegenerative tauopathies. Annu Rev Neurosci 24, 1121–1159 (2001).

3. K. A. Josephs et al., Neuropathological background of phenotypical variability in frontotemporal dementia. Acta Neuropathol 122, 137–153 (2011).

4. H. C. Tai et al., The synaptic accumulation of hyperphosphorylated tau oligomers in Alzheimer disease is associated with dysfunction of the ubiquitin-proteasome system. Am J Pathol 181, 1426–1435 (2012).

5. M. Dani et al., Microglial activation correlates in vivo with both tau and amyloid in Alzheimer’s disease. Brain 141, 2740–2754 (2018).

6. S. H. Lee, E. J. Bae, S. J. Park, S. J. Lee, Microglia-driven inflammation induces progressive tauopathies and synucleinopathies. Exp Mol Med, (2025).

7. A. Bejanin et al., Tau pathology and neurodegeneration contribute to cognitive impairment in Alzheimer’s disease. Brain 140, 3286–3300 (2017).

8. C. Langworth-Green et al., Chronic effects of inflammation on tauopathies. Lancet Neurol 22, 430–442 (2023).

9. C. Wang et al., Microglial NF-κB drives tau spreading and toxicity in a mouse model of tauopathy. Nat Commun 13, 1969 (2022).

10. Y. Kim, S. H. Ryu, J. Hyun, Y. S. Cho, Y. K. Jung, TLR2 immunotherapy suppresses neuroinflammation, tau spread, and memory loss in rTg4510 mice. Brain Behav Immun 121, 291–302 (2024).

11. D. Dutta et al., Tau fibrils induce glial inflammation and neuropathology via TLR2 in Alzheimer’s disease-related mouse models. J Clin Invest 133, (2023).

12. K. Bhaskar et al., Regulation of tau pathology by the microglial fractalkine receptor. Neuron 68, 19–31 (2010).

13. D. W. Litchfield, Protein kinase CK2: structure, regulation and role in cellular decisions of life and death. Biochem J 369, 1–15 (2003).

14. R. Gomez-Pastor et al., Abnormal degradation of the neuronal stress-protective transcription factor HSF1 in Huntington’s disease. Nat Commun 8, 14405 (2017).

15. D. Yu et al., CK2 alpha prime and alpha-synuclein pathogenic functional interaction mediates synaptic dysregulation in huntington’s disease. Acta Neuropathol Commun 10, 83 (2022).

16. Q. Zhang et al., CK2 Phosphorylating I 2 PP2A /SET Mediates Tau Pathology and Cognitive Impairment. Front Mol Neurosci 11, 146 (2018).

17. H. Yadikar et al., Screening of tau protein kinase inhibitors in a tauopathy-relevant cell-based model of tau hyperphosphorylation and oligomerization. PLoS One 15, e0224952 (2020).

18. L. Baum et al., Casein kinase II is associated with neurofibrillary tangles but is not an intrinsic component of paired helical filaments. Brain Res 573, 126–132 (1992).

19. A. C. Lim, S. Y. Tiu, Q. Li, R. Z. Qi, Direct regulation of microtubule dynamics by protein kinase CK2. J Biol Chem 279, 4433–4439 (2004).

20. R. Kimura, N. Matsuki, Protein kinase CK2 modulates synaptic plasticity by modification of synaptic NMDA receptors in the hippocampus. J Physiol 586, 3195–3206 (2008).

21. H. J. Chung, Y. H. Huang, L. F. Lau, R. L. Huganir, Regulation of the NMDA receptor complex and trafficking by activity-dependent phosphorylation of the NR2B subunit PDZ ligand. J Neurosci 24, 10248–10259 (2004).

22. S. C. Lenzken et al., Recruitment of casein kinase 2 is involved in AbetaPP processing following cholinergic stimulation. J Alzheimers Dis 20, 1133–1141 (2010).

23. M. Raftery et al., Phosphorylation of apolipoprotein-E at an atypical protein kinase CK2 PSD/E site in vitro. Biochemistry 44, 7346–7353 (2005).

24. J. Walter, A. Schindzielorz, B. Hartung, C. Haass, Phosphorylation of the beta-amyloid precursor protein at the cell surface by ectocasein kinases 1 and 2. J Biol Chem 275, 23523–23529 (2000).

25. Y. Bian et al., Global screening of CK2 kinase substrates by an integrated phosphoproteomics workflow. Sci Rep 3, 3460 (2013).

26. S. F. Rusin, M. E. Adamo, A. N. Kettenbach, Identification of Candidate Casein Kinase 2 Substrates in Mitosis by Quantitative Phosphoproteomics. Front Cell Dev Biol 5, 97 (2017).

27. B. Guerra, S. Siemer, B. Boldyreff, O. G. Issinger, Protein kinase CK2: evidence for a protein kinase CK2beta subunit fraction, devoid of the catalytic CK2alpha subunit, in mouse brain and testicles. FEBS Lett 462, 353–357 (1999).

28. M. Montenarh, C. Götz, Protein Kinase CK2α’, More than a Backup of CK2α. Cells 12, (2023).

29. I. Ceglia, M. Flajolet, H. Rebholz, Predominance of CK2α over CK2α’ in the mammalian brain. Mol Cell Biochem 356, 169–175 (2011).

30. D. Y. Lou et al., The alpha catalytic subunit of protein kinase CK2 is required for mouse embryonic development. Mol Cell Biol 28, 131–139 (2008).

31. X. Xu, P. A. Toselli, L. D. Russell, D. C. Seldin, Globozoospermia in mice lacking the casein kinase II alpha’ catalytic subunit. Nat Genet 23, 118–121 (1999).

32. D. S. Iimoto, E. Masliah, R. DeTeresa, R. D. Terry, T. Saitoh, Aberrant casein kinase II in Alzheimer’s disease. Brain Res 507, 273–280 (1990).

33. J. E. Allende, C. C. Allende, Protein kinases. 4. Protein kinase CK2: an enzyme with multiple substrates and a puzzling regulation. FASEB J 9, 313–323 (1995).

34. A. F. Rosenberger et al., Increased occurrence of protein kinase CK2 in astrocytes in Alzheimer’s disease pathology. J Neuroinflammation 13, 4 (2016).

35. M. V. Aksenova, G. S. Burbaeva, K. V. Kandror, D. V. Kapkov, A. S. Stepanov, The decreased level of casein kinase 2 in brain cortex of schizophrenic and Alzheimer’s disease patients. FEBS Lett 279, 55–57 (1991).

36. T. Saitoh, D. Iimoto, Aberrant protein phosphorylation and cytoarchitecture in Alzheimer’s disease. Prog Clin Biol Res 317, 769–780 (1989).

37. J. A. Miller et al., Neuropathological and transcriptomic characteristics of the aged brain. Elife 6, (2017).

38. Y. Yoshiyama et al., Synapse loss and microglial activation precede tangles in a P301S tauopathy mouse model. Neuron 53, 337–351 (2007).

39. C. K. Walker et al., Dendritic Spine Remodeling and Synaptic Tau Levels in PS19 Tauopathy Mice. Neuroscience 455, 195–211 (2021).

40. J. P. Anderson et al., Phosphorylation of Ser-129 is the dominant pathological modification of alpha-synuclein in familial and sporadic Lewy body disease. J Biol Chem 281, 29739–29752 (2006).

41. H. Takeuchi et al., P301S mutant human tau transgenic mice manifest early symptoms of human tauopathies with dementia and altered sensorimotor gating. PLoS One 6, e21050 (2011).

42. Y. Zhou et al., The Alzheimer’s Cell Atlas (TACA): A single-cell molecular map for translational therapeutics accelerator in Alzheimer’s disease. Alzheimers Dement (N Y) 8, e12350 (2022).

43. E. Gerrits et al., Distinct amyloid-β and tau-associated microglia profiles in Alzheimer’s disease. Acta Neuropathol 141, 681–696 (2021).

44. M. Otero-Garcia et al., Molecular signatures underlying neurofibrillary tangle susceptibility in Alzheimer’s disease. Neuron 110, 2929–2948.e2928 (2022).

45. K. Leng et al., Molecular characterization of selectively vulnerable neurons in Alzheimer’s disease. Nat Neurosci 24, 276–287 (2021).

46. S. F. Lau, H. Cao, A. K. Y. Fu, N. Y. Ip, Single-nucleus transcriptome analysis reveals dysregulation of angiogenic endothelial cells and neuroprotective glia in Alzheimer’s disease. Proc Natl Acad Sci U S A 117, 25800–25809 (2020).

47. C. A. Marshall et al., Inhibition of CK2 mitigates Alzheimer’s tau pathology by preventing NR2B synaptic mislocalization. Acta Neuropathol Commun 10, 30 (2022).

48. B. R. Hoover et al., Tau mislocalization to dendritic spines mediates synaptic dysfunction independently of neurodegeneration. Neuron 68, 1067–1081 (2010).

49. Y. Shi et al., Microglia drive APOE-dependent neurodegeneration in a tauopathy mouse model. J Exp Med 216, 2546–2561 (2019).

50. Y. Shi et al., ApoE4 markedly exacerbates tau-mediated neurodegeneration in a mouse model of tauopathy. Nature 549, 523–527 (2017).

51. H. Martini-Stoica et al., TFEB enhances astroglial uptake of extracellular tau species and reduces tau spreading. J Exp Med 215, 2355–2377 (2018).

52. H. Takahashi et al., Reduced progranulin increases tau and α-synuclein inclusions and alters mouse tauopathy phenotypes via glucocerebrosidase. Nat Commun 15, 1434 (2024).

53. A. Litvinchuk et al., Complement C3aR Inactivation Attenuates Tau Pathology and Reverses an Immune Network Deregulated in Tauopathy Models and Alzheimer’s Disease. Neuron 100, 1337–1353.e1335 (2018).

54. J. Kim et al., Cerebral transcriptome analysis reveals age-dependent progression of neuroinflammation in P301S mutant tau transgenic male mice. Brain Behav Immun 80, 344–357 (2019).

55. F. H. Shiu et al., ADGRB1 contributes to astrocyte-mediated phagocytosis of excitatory synapses. Exp Neurol 395, 115451 (2025).

56. T. Lala, R. A. Hall, Adhesion G protein-coupled receptors: structure, signaling, physiology, and pathophysiology. Physiol Rev 102, 1587–1624 (2022).

57. X. Chen et al., Microglia-mediated T cell infiltration drives neurodegeneration in tauopathy. Nature 615, 668–677 (2023).

58. K. F. Odfalk, K. F. Bieniek, S. C. Hopp, Microglia: Friend and foe in tauopathy. Prog Neurobiol 216, 102306 (2022).

59. C. Gao, J. Jiang, Y. Tan, S. Chen, Microglia in neurodegenerative diseases: mechanism and potential therapeutic targets. Signal Transduct Target Ther 8, 359 (2023).

60. S. Fixemer et al., Microglia phenotypes are associated with subregional patterns of concomitant tau, amyloid-β and α-synuclein pathologies in the hippocampus of patients with Alzheimer’s disease and dementia with Lewy bodies. Acta Neuropathol Commun 10, 36 (2022).

61. R. Hassan-Abdi, A. Brenet, M. Bennis, C. Yanicostas, N. Soussi-Yanicostas, Neurons Expressing Pathological Tau Protein Trigger Dramatic Changes in Microglial Morphology and Dynamics. Front Neurosci 13, 1199 (2019).

62. S. Hong et al., Complement and microglia mediate early synapse loss in Alzheimer mouse models. Science 352, 712–716 (2016).

63. B. Dejanovic et al., Changes in the Synaptic Proteome in Tauopathy and Rescue of Tau-Induced Synapse Loss by C1q Antibodies. Neuron 100, 1322–1336.e1327 (2018).

64. H. Lui et al., Progranulin Deficiency Promotes Circuit-Specific Synaptic Pruning by Microglia via Complement Activation. Cell 165, 921–935 (2016).

65. K. Pampuscenko et al., Extracellular tau stimulates phagocytosis of living neurons by activated microglia via Toll-like 4 receptor-NLRP3 inflammasome-caspase-1 signalling axis. Sci Rep 13, 10813 (2023).

66. J. Leyh et al., Classification of Microglial Morphological Phenotypes Using Machine Learning. Front Cell Neurosci 15, 701673 (2021).

67. S. Parhizkar et al., Lemborexant ameliorates tau-mediated sleep loss and neurodegeneration in males in a mouse model of tauopathy. Nat Neurosci, (2025).

68. E. R. Roy et al., Concerted type I interferon signaling in microglia and neural cells promotes memory impairment associated with amyloid β plaques. Immunity 55, 879–894.e876 (2022).

69. Y. B. Choi et al., Erythropoietin-derived peptide treatment reduced neurological deficit and neuropathological changes in a mouse model of tauopathy. Alzheimers Res Ther 13, 32 (2021).

70. S. L. DeVos et al., Tau reduction prevents neuronal loss and reverses pathological tau deposition and seeding in mice with tauopathy. Sci Transl Med 9, (2017).

71. J. Wagner et al., Reducing tau aggregates with anle138b delays disease progression in a mouse model of tauopathies. Acta Neuropathol 130, 619–631 (2015).

72. N. Zheng et al., Electrophysiology-based screening identifies neuronal HtrA serine peptidase 2 (HTRA2) as a synaptic plasticity regulator participating in tauopathy. Transl Psychiatry 15, 5 (2025).

73. R. El Fatimy et al., MicroRNA-132 provides neuroprotection for tauopathies via multiple signaling pathways. Acta Neuropathol 136, 537–555 (2018).

74. T. Illouz et al., Unbiased classification of spatial strategies in the Barnes maze. Bioinformatics 32, 3314–3320 (2016).

75. K. Chen et al., Selective removal of astrocytic PERK protects against glymphatic impairment and decreases toxic aggregation of β-amyloid and tau. Neuron, (2025).

76. I. I. N. Da Silva et al., CK2 inhibition suppresses glial inflammation in the brain. bioRxiv, (2025).

77. M. Pérez, J. Avila, The expression of casein kinase 2alpha’ and phosphatase 2A activity. Biochim Biophys Acta 1449, 150–156 (1999).

78. J. Liu, J. Tian, R. Xie, L. Chen, CK2 inhibitor DMAT ameliorates spinal cord injury by increasing autophagy and inducing anti-inflammatory microglial polarization. Neurosci Lett 805, 137222 (2023).

79. S. Mishra, C. Kinoshita, A. D. Axtman, J. E. Young, Evaluation of a Selective Chemical Probe Validates That CK2 Mediates Neuroinflammation in a Human Induced Pluripotent Stem Cell-Derived Microglial Model. Front Mol Neurosci 15, 824956 (2022).

80. I. Y. Chen et al., Upregulation of the chemokine (C-C motif) ligand 2 via a severe acute respiratory syndrome coronavirus spike-ACE2 signaling pathway. J Virol 84, 7703–7712 (2010).

81. S. R. Larson et al., Myeloid Cell CK2 Regulates Inflammation and Resistance to Bacterial Infection. Front Immunol 11, 590266 (2020).

82. D. Drygin et al., Protein kinase CK2 modulates IL-6 expression in inflammatory breast cancer. Biochem Biophys Res Commun 415, 163–167 (2011).

83. K. Parhar, J. Morse, B. Salh, The role of protein kinase CK2 in intestinal epithelial cell inflammatory signaling. Int J Colorectal Dis 22, 601–609 (2007).

84. M. Yamada et al., Inhibition of protein kinase CK2 prevents the progression of glomerulonephritis. Proc Natl Acad Sci U S A 102, 7736–7741 (2005).

85. J. Huang et al., Sphingosine kinase 1 mediates diabetic renal fibrosis via NF-κB signaling pathway: involvement of CK2α. Oncotarget 8, 88988–89004 (2017).

86. N. N. Singh, D. P. Ramji, Protein kinase CK2, an important regulator of the inflammatory response? J Mol Med (Berl) 86, 887–897 (2008).

87. J. Avila, J. J. Lucas, M. Perez, F. Hernandez, Role of tau protein in both physiological and pathological conditions. Physiol Rev 84, 361–384 (2004).

88. N. Zarate et al., Neurochemical correlates of synapse density in a Huntington’s disease mouse model. J Neurochem 164, 226–241 (2023).

89. E. M. Coomans et al., In vivo tau pathology is associated with synaptic loss and altered synaptic function. Alzheimers Res Ther 13, 35 (2021).

90. T. Vogels, A. N. Murgoci, T. Hromádka, Intersection of pathological tau and microglia at the synapse. Acta Neuropathol Commun 7, 109 (2019).

91. X. Liu et al., Proteomic analysis of ferroptosis pathways reveals a role of CEPT1 in suppressing ferroptosis. Protein Cell 15, 686–703 (2024).

92. P. Yin et al., Aged monkey brains reveal the role of ubiquitin-conjugating enzyme UBE2N in the synaptosomal accumulation of mutant huntingtin. Hum Mol Genet 24, 1350–1362 (2015).

93. J. Y. Qiu et al., OXCT1 regulates hippocampal neurogenesis and alleviates cognitive impairment via the Akt/GSK-3β/β-catenin pathway after subarachnoid hemorrhage. Brain Res 1827, 148758 (2024).

94. J. M. Serfass et al., Endophilin B2 facilitates endosome maturation in response to growth factor stimulation, autophagy induction, and influenza A virus infection. J Biol Chem 292, 10097–10111 (2017).

95. Y. H. Wang et al., Endophilin B2 promotes inner mitochondrial membrane degradation by forming heterodimers with Endophilin B1 during mitophagy. Sci Rep 6, 25153 (2016).

96. O. Bohana-Kashtan, L. A. Pinna, Z. Fishelson, Extracellular phosphorylation of C9 by protein kinase CK2 regulates complement-mediated lysis. Eur J Immunol 35, 1939–1948 (2005).

97. S. Pelloux et al., Identification of a cryptic protein kinase CK2 phosphorylation site in human complement protease Clr, and its use to probe intramolecular interaction. FEBS Lett 386, 15–20 (1996).

98. D. Mudaliar et al., Discovery of a CK2α’-Biased ATP-Competitive Inhibitor from a High-Throughput Screen of an Allosteric-Inhibitor-Like Compound Library. ACS Chem Neurosci, (2024).

99. S. C. Vermilyea et al., Loss of tau expression attenuates neurodegeneration associated with α-synucleinopathy. Transl Neurodegener 11, 34 (2022).

100. T. G. Brown et al., Striatal spatial heterogeneity, clustering, and white matter association of GFAP. Front Cell Neurosci 17, 1094503 (2023).

101. M. A. Solem et al., Absence of hippocampal pathology persists in the Q175DN mouse model of Huntington’s disease despite elevated HTT aggregation. J Huntingtons Dis 14, 59–84 (2025).

102. P. Bankhead et al., QuPath: Open source software for digital pathology image analysis. Sci Rep 7, 16878 (2017).

103. A. C. Ruifrok, D. A. Johnston, Quantification of histochemical staining by color deconvolution. Anal Quant Cytol Histol 23, 291–299 (2001).

104. J. Schindelin et al., Fiji: an open-source platform for biological-image analysis. Nat Methods 9, 676–682 (2012).

105. K. Young, H. Morrison, Quantifying Microglia Morphology from Photomicrographs of Immunohistochemistry Prepared Tissue Using ImageJ. J Vis Exp, (2018).

106. I. Arganda-Carreras, R. Fernández-González, A. Muñoz-Barrutia, C. Ortiz-De-Solorzano, 3D reconstruction of histological sections: Application to mammary gland tissue. Microsc Res Tech 73, 1019–1029 (2010).

107. N. Zarate et al., Heat Shock Factor 1 Directly Regulates Postsynaptic Scaffolding PSD-95 in Aging and Huntington’s Disease and Influences Striatal Synaptic Density. Int J Mol Sci 22, (2021).

108. S. U. McKinstry et al., Huntingtin is required for normal excitatory synapse development in cortical and striatal circuits. J Neurosci 34, 9455–9472 (2014).

109. M. W. Pitts, Barnes Maze Procedure for Spatial Learning and Memory in Mice. Bio Protoc 8, (2018).

110. C. A. Barnes, Memory deficits associated with senescence: a neurophysiological and behavioral study in the rat. J Comp Physiol Psychol 93, 74–104 (1979).

111. M. I. Love, W. Huber, S. Anders, Moderated estimation of fold change and dispersion for RNA-seq data with DESeq2. Genome Biol 15, 550 (2014).

112. U. Raudvere et al., g:Profiler: a web server for functional enrichment analysis and conversions of gene lists (2019 update). Nucleic Acids Res 47, W191–W198 (2019).

