## Supplementary Methods and Figures for "Protein kinase CK2α’ as a dual modulator of immune signaling and synaptic dysfunction in Tauopathy"

**Supplemental methods and materials**

**Primary cell culture**

Primary cell culture was performed as previously described *(99)* . Briefly the hippocampus was dissected from mouse pups between postnatal day 0-2, placed in cold Hibernate-A medium (BrainBits LLC; Springfield, Illinois, USA), then transferred to digestion medium (Hibernate-A -CaCl2 (BrainBits, Cordova, TN), papain, L-Cysteine, and EDTA) for 15 minutes minutes at 37°C with occasional gentle shaking. Hippocampi were then washed in a digestion inhibitor solution before being triturated with a pasteur pipette. Cells were plated in 35mm cell culture dish at a density of 1x10^6^ cells/plate and placed in 37°C/5% CO2 incubator. The next day media was replaced with NbActiv4 (BrainBits, Cordova, TN) +FdU mitotic inhibitor.

For RNA collection cells were transfected at 14 days in vitro (DIV) with siCK2α’ or ScrRNA as described in cell culturing techniques, RNA was extracted 48 hours later as described. For spine imaging cells were transfected with a plasmid expressing DsRed (Clonetech; Mountain View, CA) and Tau-P301S-EGFP under the CMV promotor in the pRK5 vector at 9 DIV (LTX DNA transfection kit (#15338030, Thermo Fisher Scientific; Waltham, MA, USA) and then transfected with siCK2α’ or ScrRNA at 21DIV. 48 hours after siRNA transfection (23 DIV) cells were live imaged using Leica Stellaris 8 confocal microscope with gas and temperature-regulated culture dish chamber. Z-stacks were obtained at 63x x 2.5 magnification. Leica software "Lightening" noise reduction was applied to the z-stack series and then maximum projection image was obtained for subsequent analysis.

Dendrite density analysis was performed by blinded experimenter. For each image spines were counted in two different 20μm long dendrite regions. An average dendritic density was then calculated and averaged per image and relativized to control.

**RNA extraction and qPCR**

RNA was extracted from cells and mouse striatal tissues using the RNeasy extraction kit (Qiagen) and reverse-transcribe with Superscript First Strand Synthesis System (Invitrogen). SYBR green based PCR was performed with SYBR mix (Roche) using the LightCycler 480 System (Roche). Samples were run in triplicate and normalized to GAPDH.

For qPCR the following primers were used: CK2α’ **FWD-**CGACTGATTGATTGGGGTCT **REV-**AGAATGGCTCCTTTCGGAAT; CK2α **FWD-**TCCCCATGCTGTGACAATAA **REV-**AAGACCCTGTGTCACGAACC; CK2β **FWD-** AGTCCTCCAGACACCACCAC **REV-**GACTGGGCTCTTGAAGTTGC; huTau **FWD-** GCTGGCCTGAAAGCTGAAGA **REV-** CGTTTTACCATCAGCCCCCT.

**Microglia morphology**

Microglia morphology was quantified as described *(105).* Iba1-stained slices were imaged using a Lecia Stellaris confocal microscope at 40x (2.5xzoom) z-stacks were acquired at a 0.3μm step size. Nine cells per animal (3 slices x 3 cells/slice) were analyzed. Maximum projection were processed in FIJI: brightness/contrast adjust, converted to binary, and despeckled to remove noise and then skeletonized. The AnalyzeSkeleton plugin *(106)* was used for quantification.

**Cytokine proteome profiling**

Cytokine and chemokine levels were assessed using the Proteome Profiler Mouse Cytokine Array Panel (ARY006, R&D Systems) following manufacturer instructions. Frozen hippocampi from 9-10 months old PS19 and PS19;CK2α’^(+/-)^ mice (n=genotype;2F/2M) were homogenized in PBS with Halt protease inhibitor cocktail/phosphatase inhibitors (Fisher Scientific) and 1% triton X-100 (Sigma). Lysates were frozen at -80°C for 15 min, thawed and centrifuged at 10,000 × g for 5 min. 300 μg of protein were applied per membrane. Imaging was conducted as described for immunoblotting. Spot intensity was quantified using FIJI software and a batch correction was applied. Membrane spots corresponding to 40 cytokines or chemokines were measured as outlined in manufacturer protocol.

***in situ* hybridization**

Tissues were collected as previously described in the immunohistochemistry procedure. Slices were mounted stored at -80ºC overnight before *in situ* hybridization. RNAscope was performed according to the ACDbio manufacturer’s protocol using a custom probe for *Csnk2a2*. Immediately following RNAScope protocol, the slides were blocked with 10% Normal Goat Serum (NGS) in Co-detection antibody diluent (CDD) for 1 hour. Anti-IBA1 (Rabbit, Fujifilm Wako 019-19741) was diluted at 1:200 in 5% NGS with CDD and slides were incubated overnight. Slides were incubated the next day in Alexa-fluorophore-conjugated antibodies (Invitrogen) (1:40 in CDD with 5% NGS) for 2h at room temperature. Slides were mounted in ProLong Gold Antifade with DAPI (Invitrogen) and subsequently imaged.

**Tau pathology type**

Tau pathology type analysis was conducted by 3 independent and blinded investigators. Tau pathologies were defined based on previous publications *(49-52)* and based on the pattern of AT8+ staining. Briefly as described by Shi et al 2017 *(50)* type 1; displays mossy fiber and sparse and diffuse cell body staining in the dentate gyrus (DG), type 2; displays compact dense tangle like cell body staining in the DG and CA3 with some sparse CA1 cell body staining, type 3; staining is seen in the neuropil of the stratum radiatum of the CA1 staining along with staining of dendrites from pyramidal neurons and sparse cell body staining, type 4; dense fragmented, dotted, and grainy staining is observed all over the hippocampus. The percentage of pathology for the cohort for each type was determined by the following calculation; ((number of mice with pathology type)/(total number of mice in cohort)) x 100.

**Image quantification**

NeuN quantifications were done using QuPath cell detection. Identical ROIs were applied to each image for all 3 regions of the hippocampus. The cell detection tool was used to automatically detect NeuN+ cells based on identical thresholding settings applied to all images.

Quantification of Iba1+ and GFAP+ cell was conducted as previously described *(100,101)*. Average projections of 20x tiled z-stack confocal image were generated in FIJI for analysis. Two ROIs (~3900 um^2^) were drawn within the CA1, CA3, and DG and cells were manually counted using FIJI cell counter. Cells were classified as Iba1+ or GFAP+ when DAPI nuclei overlapped with the stained area. Counts were averaged and normalized per mm^2^ and reported as animal averages (2 ROIs/slice; 3 slices/animal).

Quantification of DAB positive percent area was quantified using either the Colordevonvolution2 plugin in FIJI *(103)* or in QuPath. Briefly, a threshold of positive DAB staining was determined and applied to all images to then calculate the percentage positive area.

Quantification of PSD95+ puncta within Iba1+ microglia were performed using Imaris software (Bitplane). A confocal microscope (Stellaris, Lecia) was used to acquire images in the CA1 stratum radiatum region of the hippocampus with a 40x objective and 3x zoom factor with 0.34 size z-steps for a total of 15 steps. Lecia Stellaris software lighting processing was applied before analysis. Images were then 3D rendered in Imaris and cropped to isolate individual Iba1+ cells (2 per slice). Iba1+ surfaces were generated and used to mask PSD95+ signal. The masked PSD95+ signal was then used to create spots, to ensure it was located within Iba1+ signal. The number of spots within the Iba1+ surfaces per cell soma defined as the region where Iba1+ signal colocalized with DAPI and within the whole Iba1+ cell surface were counted and reported as a number.

**Supplemental Figures**

**Figure S1**

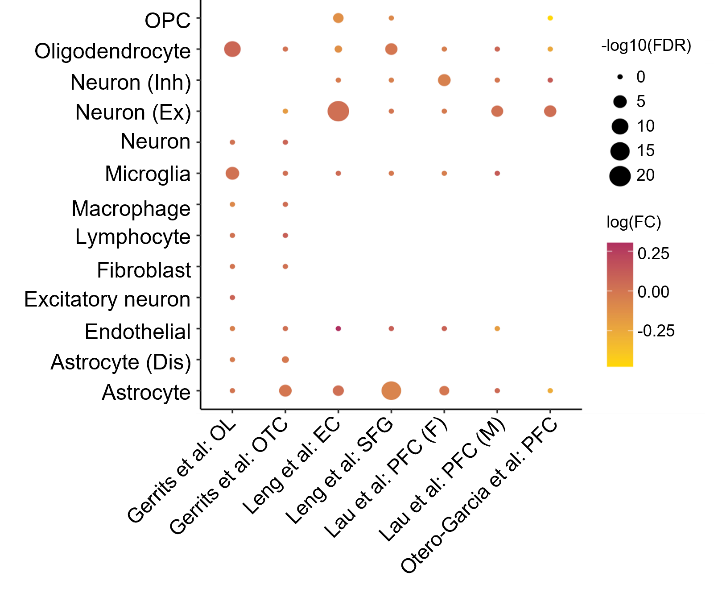

**Figure S1. *CSNK2a2* scRNA-seq expression in human brain tissue.** **(A)** Summary of *CSNK2a2* single cell expression from studies found in The Alzheimer's Cell Atlas (TACA) in occipital lobe (OL), occipitotemporal cortex (OTC), entorhinal cortex (EC), superior frontal gyrus (SFG), and prefrontal cortex (PFC). Absence of expression for a cell type indicates that cell type was not identified as a cluster in the study.

**Figure S2**

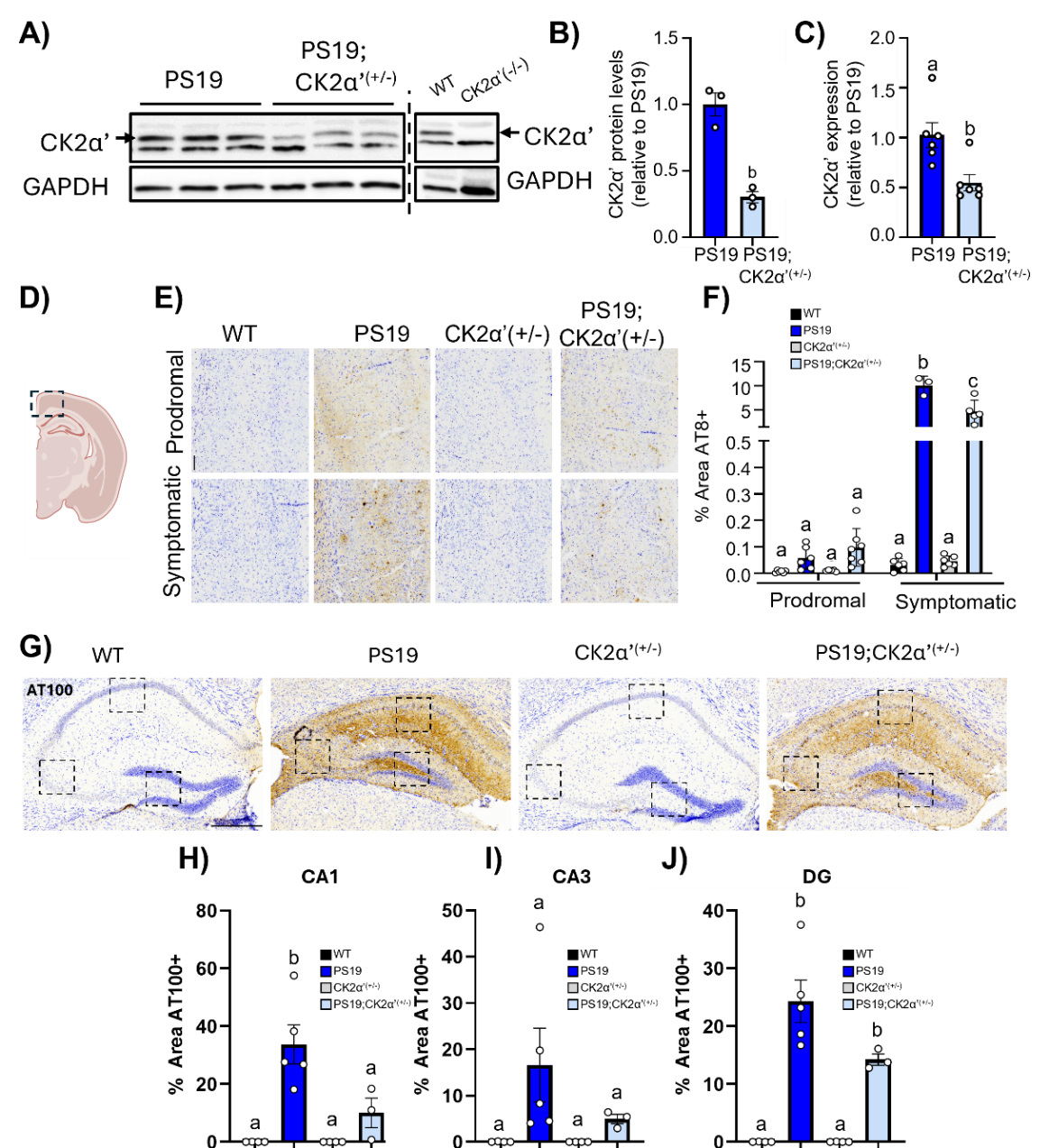

**Figure S2. PS19;CK2α’(+/-) mice display decreased AT8 staining in the cortex and AT100 in hippocampus of symptomatic mice. (A)** Representative western blot of CK2α’ in 7m frontal cortex of PS19 animals. **(B)** Quantifications of CK2α’ protein levels quantified via ImageJ from immunoblot. **(C)** RT-qPCR results for CK2α’ gene expression levels. **(D)** Representative diagram of cortical region imaged for quantification in E and F. **(E)** Representative images of AT8 staining in the cortex of symptomatic mice. **(F)** Quantification of % area AT8+ in the cortex of prodromal and symptomatic mice (n=3-7 mice/genotype). **(G)** Representative images of AT100 staining in the hippocampus of symptomatic mice. **(H-J)** Quantification of % area AT100+ in the hippocampus of symptomatic mice (n=3-5/genotype) Data are shown as mean ± SEM. Statistical analyses were conducted using a unpaired t-test (B,C), Two-way ANOVA (F) or One-way ANOVA (H-J) with Tukey’s post-hoc analysis ( and displayed with compact letter display. Panel D was created in BioRender (White, A. (2025) <https://BioRender.com/eqznm6m>).

**Figure S3**

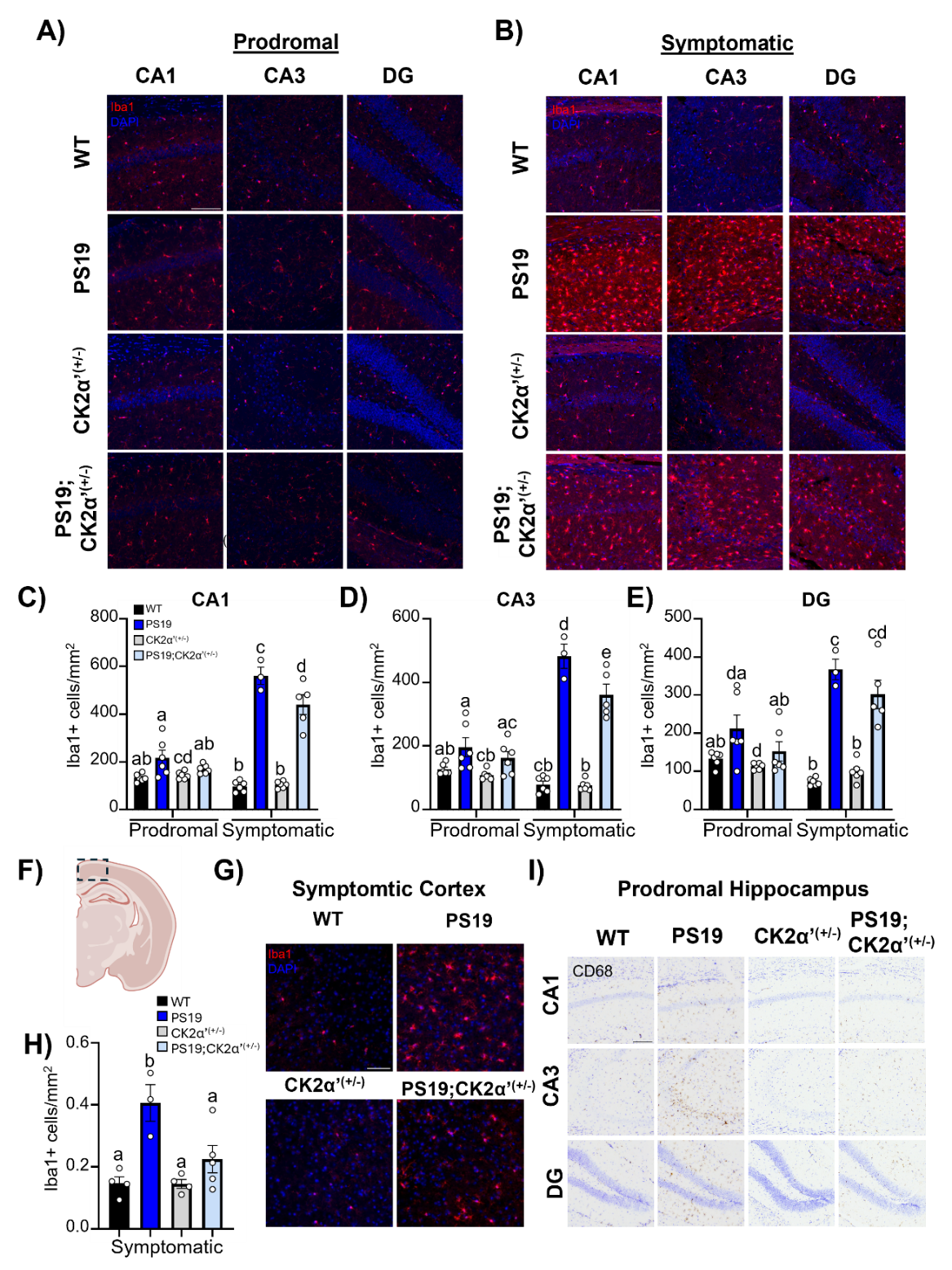

**Figure S3. CK2α’ haploinsufficency decreased Iba1+ microglia cell number in the hippocampus and overlaying cortex of PS19 mice. (A-B)** Immunostaining for Iba1 in the CA1, CA3 and DG of the hippocampus in prodromal (**A**) and symptomatic (**B**) cohorts. Scale bar=100 μm. **(C-E)** Quantification of Iba1+ cell counts in CA1, CA3 and DG of hippocampus (n=3-6 mice/genotype). **(F)** Representative area corresponding to overlying cortex is indicated with a boxed area. **(G)** Representative Iba1 immunostaining in the cortex scale bar=50 μm. **(H)** Quantification of Iba1+ cells in the cortex (n=3-6 mice/genotype). All data are shown as mean ± SEM. Statistical analyses were conducted using two-way ANOVA with tukey’s multiple corrections and represented with compact letter display. Panel **F** was created in BioRender (White, A. (2025) <https://BioRender.com/eqznm6m>).

**Figure S4**

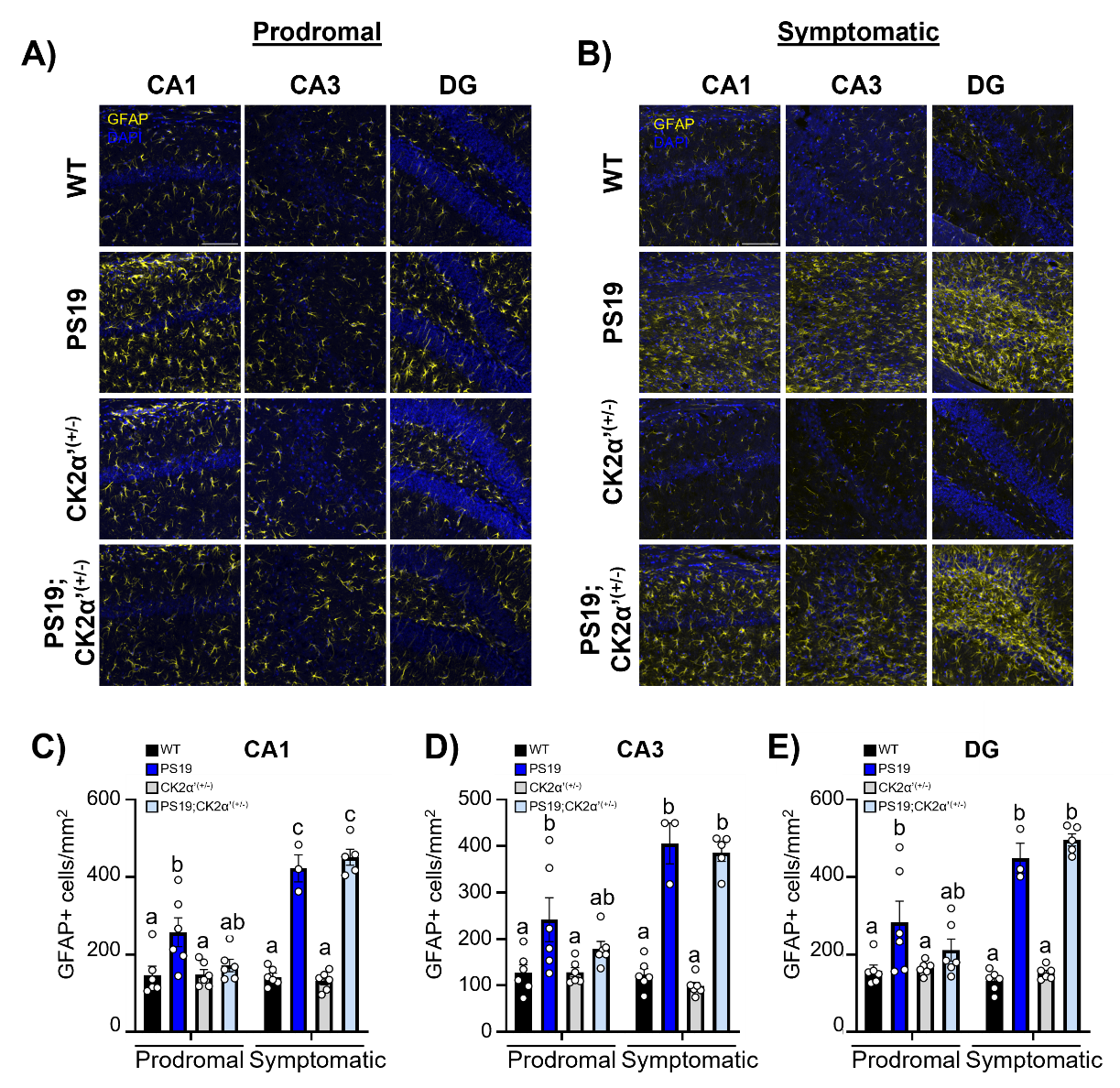

**Figure S4. CK2α’ haploinsufficency decreased GFAP+ astrocytes in the overlaying cortex but not in the hippocampus of PS19 mice. (A-B)** Immunostaining for GFAP in the CA1, CA3 and DG of the hippocampus in prodromal (A) and symptomatic (B) cohorts. Scale bar=100 μm. **(C-E)** Quantification of GFAP+ cells in CA1, CA3 and DG of hippocampus. (n=3-6 mice/genotype). **(F)** Representative area corresponding to overlying cortex is indicated with a boxed area. Statistical analyses were conducted using two-way ANOVA with tukey’s multiple corrections and represented with compact letter display. Panel **F** was created in BioRender (White, A. (2025) <https://BioRender.com/eqznm6m>).

**Figure S5**

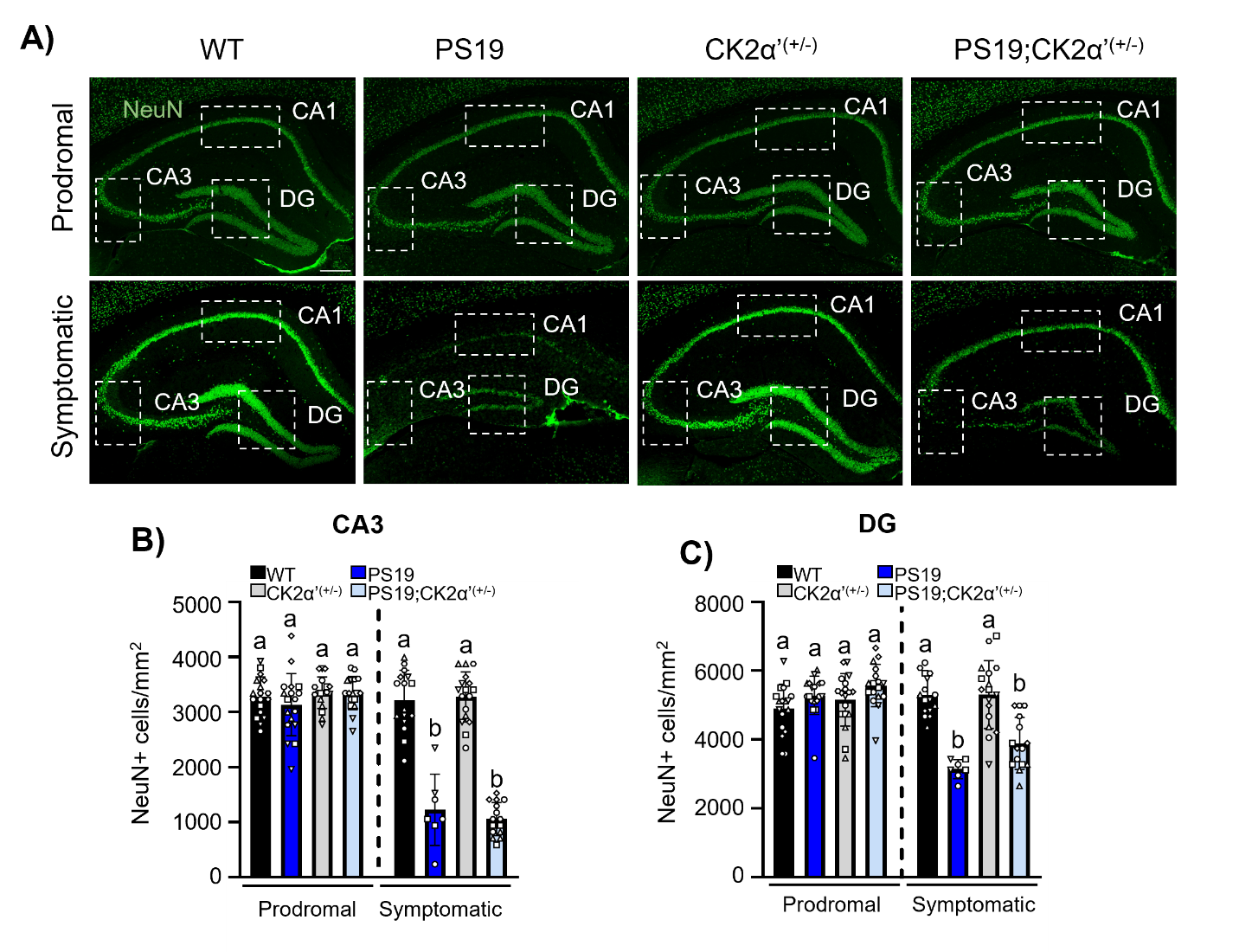

**Supplementary Figure 5. NeuN pathology and variability among symptomatic PS19 mice. (A)** NeuN immunostaining in the hippocampus of prodromal and symptomatic mice. Dotted boxes indicate quantified subregions: CA1, CA3, and DG. Scale bar = 250 μm. **(B-D)** Quantifications of NeuN+ cells (cells/mm^2^) in the CA3 (**B**), and DG (**C**) in prodromal and symptomatic cohorts (n= 3-6 mice/genotype;2-3 slices/mouse;1 data point=1 slice, shape of data point indicates slices of the same animal). Data was shown as mean ± SEM. Statistical analyses were conducted using one way-ANOVA with Tukey’s post-hoc analysis and displayed by compact letter display. Comparisons shown relative to cohort age group.

**Figure S6**

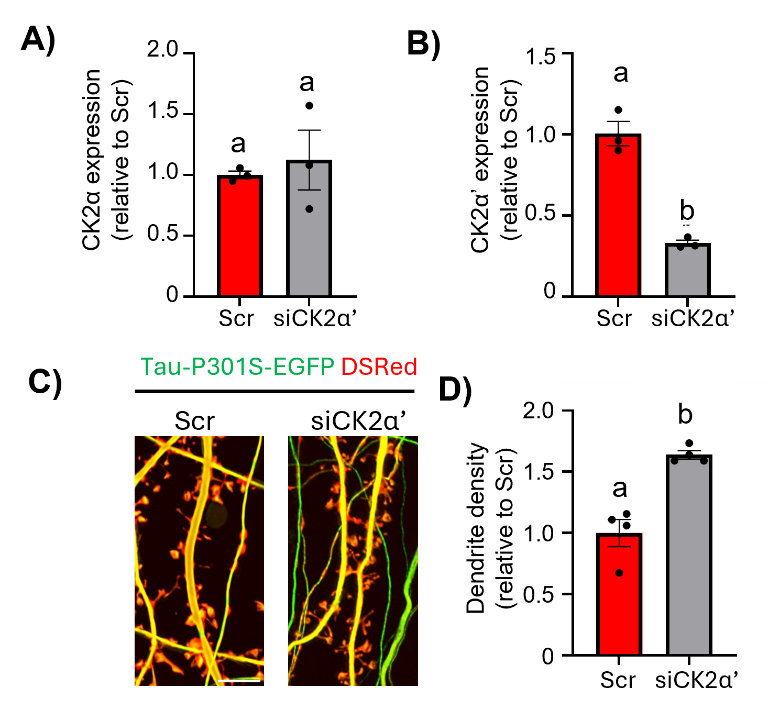

**Supplementary Figure 6. Silencing CK2α’ in primary cortical cells expressing Tau-P301L increases dendritic spine density. (A-B)** RT-qPCR analysis of mRNA expression levels of CK2α’ and CK2α (n=3/group). **(C)** Representative images of primary culture with DS-red and Tau-P301L-EGFP, scale bar=5um. **(D)** Quantification of spine density per 20um dendrite region shown relative to Scr. Data shown as mean ± SEM, significance determined by unpaired t-test and displayed with compact letter display

**Figure S7**

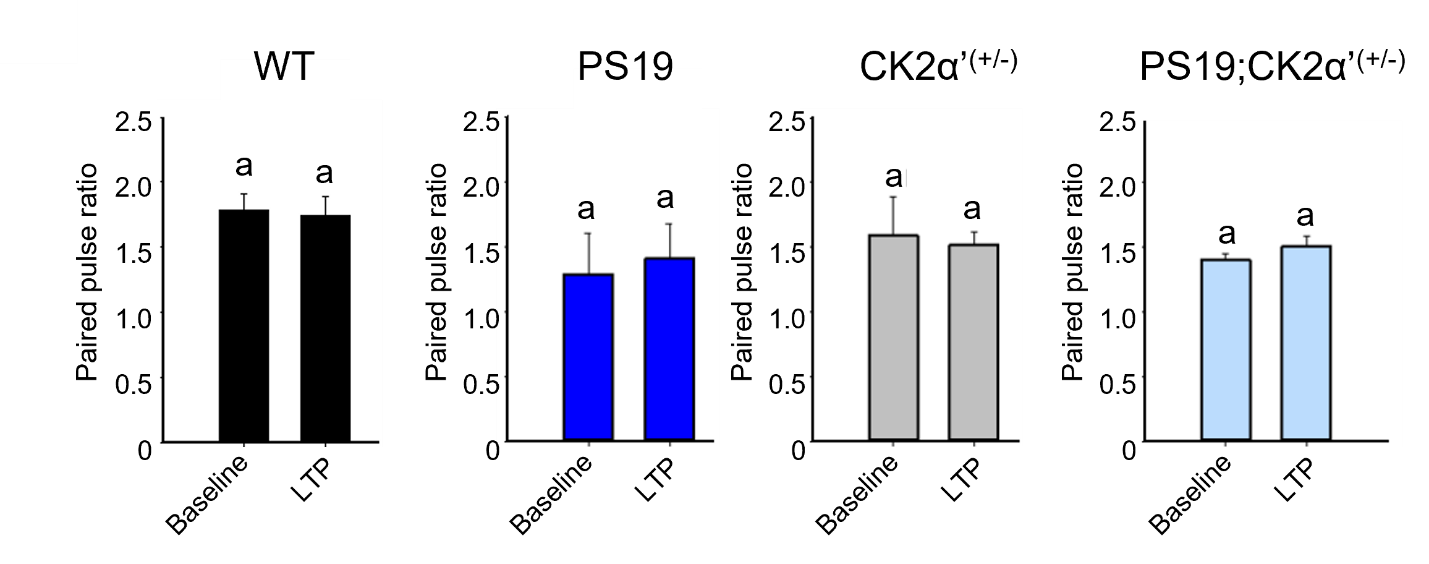

**Supplementary Figure 7. Paired pulse ratio of hippocampal slices shows no deficits between genotypes. (A)** Paired-pulse ratio (PPR) in acute hippocampal slices of (WT, Black n=5; CK2α’^(+/-)^, Grey, n=5; PS19, Dark Blue, n=5; PS19; CK2α’^(+/-)^, Light blue n=4). Data are shown as mean ± SEM. Statistical analyses were conducted using two-sided Student’s t test Significance displayed with compact letter display.

**Figure S8**

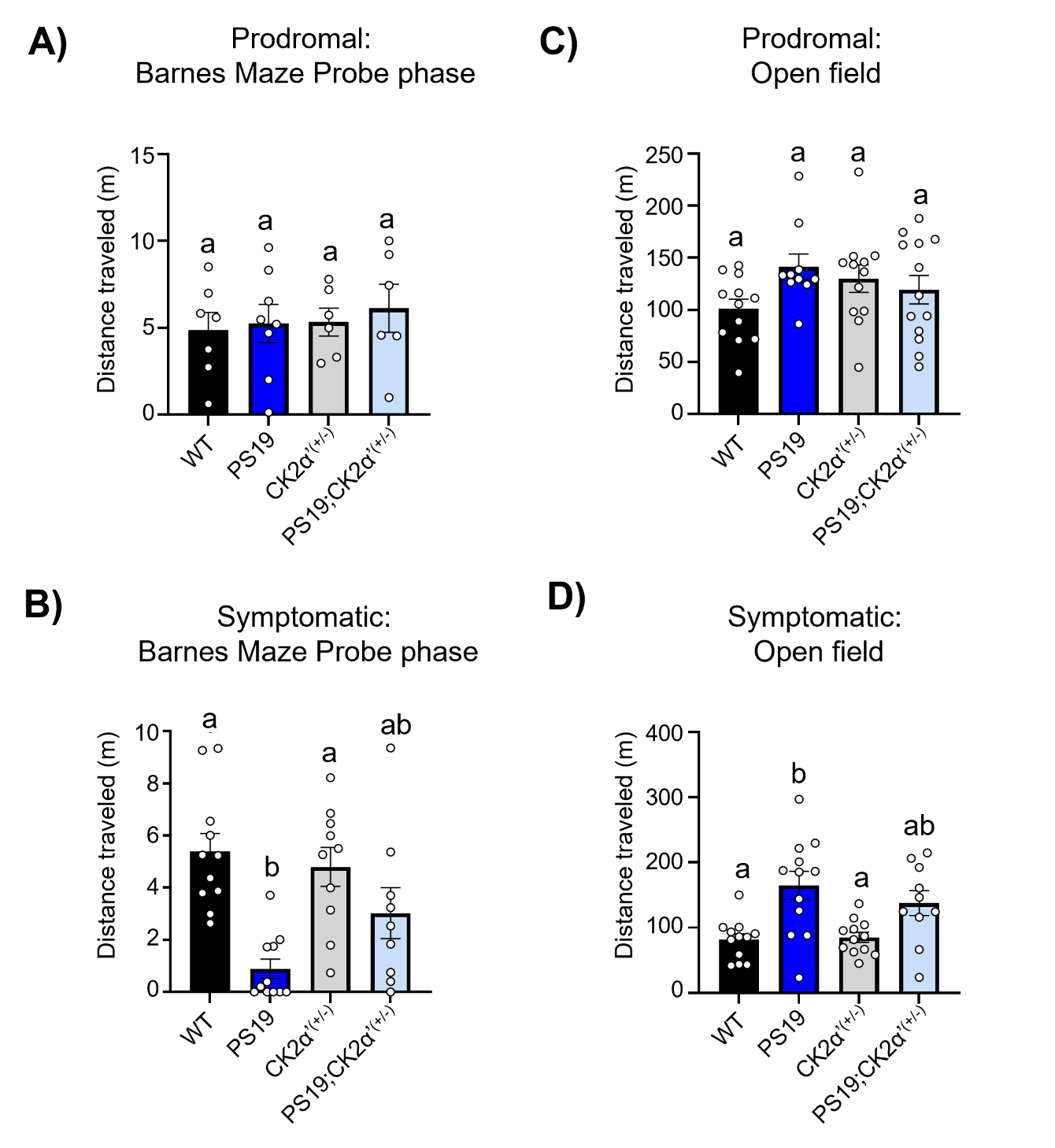

**Supplementary Figure 8. Symptomatic PS19 demonstrates decreased movement on Barnes Maze, but hyperactive phenotype in open field. (A-B)** Distance traveled in prodromal and symptomatic mice during the probe phase of Barnes Maze. **(C-D)** Distance traveled by prodromal and symptomatic mice during 1 hour in an open field arena. Data are shown as mean ± SEM (n=10-12 mice/genotype, outliers removed using ROUT (Q=1%). Statistical analyses were conducted using One-Way ANOVA with Tukey’s post-hoc analysis. Significance displayed with compact letter display.
